# Future Sequon Finder - A novel approach for predicting future N-linked glycosylation sequons on viral surface proteins

**DOI:** 10.1101/2024.11.25.625313

**Authors:** Shane P. Bryan, Martin S. Zand

## Abstract

Influenza viruses are known to evade host immune responses by shielding vulnerable surface protein epitopes via N-linked glycosylation. A program titled *Future Sequon Finder* was developed to predict the locations in which glycan binding sites are most likely to emerge in future influenza hemagglutinin proteins. The predictive modeling approach considers how closely sites in currently circulating strains resemble glycosylation sequons genetically, the surface accessibility of those sites, and the site-specific mutation frequency of the amino acids within those sites that would need to change to generate a glycosylation sequon. The efficacy of this model is tested using historic human H1N1 and H3N2 influenza strains along with swine H1N1 strains. Through this analysis, it is revealed that glycosylation addition events in influenza hemagglutinin proteins are almost always the result of single nucleotide mutation events. It is also revealed that site-specific mutation frequency and surface accessibility are powerful predictors of which sites will become glycosylated in human influenza viruses when considered with the genetic composition of the sites in question. Having been designed to incorporate these factors, the program successfully predicted almost every historic sequon addition event and, in the case of the human strains, ranked them highly among falsely predicted sequons. After demonstrating the model’s power with historical data, the program is used to predict future HA glycosylation sequon locations based on currently circulating human influenza viruses.

## Introduction

Glycosylation of viral proteins that bind to cell surface ligands is a key evolutionary adaptation, as it alters protein antigenicity and shields target sites from immune responses [1–6]. Knowledge of actual or potential N-linked glycosylation sites is important for identifying epitopes on viral surface proteins that serve as ideal target sites for human antibodies [7], vaccine engineering [8–10], or immune epitope focusing [11–13]. The highly conserved process of N-linked glycosylation involves the attachment of a polysaccharide to an asparagine (N) residue facilitated by oligosaccharyltransferase, which recognizes a specific amino acid (AA) sequence known as a glycosylation sequon [14]. A glycosylation sequon is defined by the AA sequence ‘N-!P-S/T’ where the first AA is N, the second AA is anything but proline (P), and the third AA is either serine (S) or threonine (T) [15]. Viruses have exploited protein glycosylation mechanisms in host cells to modify their proteins, aiding in viral protein folding, transportation, host receptor binding, and antibody evasion [16]. N-linked glycan trees pivot around their N residue anchor, effectively shielding a large protein surface area and sterically blocking antibodies attempting to bind to nearby epitopes [17, 18]. This effect, known as ‘glycan shielding’, must be considered when developing monoclonal antibody therapies, vaccination strategies, and antigenic escape predictions.

Influenza hemagglutinin (HA) proteins enable the virus to bind and enter host cells via interaction with sialic acid receptors and are a key target for antibody-mediated immune responses in the host [19]. As influenza virus (IFV) strains circulate, human populations develop immune responses capable of neutralizing closely related IFVs via HA interference, resulting in population immunity that slows the spread of infection [20]. To circumvent this population immunity, IFVs are constantly modifying their HA proteins [21–24]. These modifications include increases in receptor binding avidity, distal mutations altering HA conformation and epitope accessibility, mutations in epitopes directly altering their structure, and mutations resulting in sequon addition events (SAEs) that allow the binding of new glycans capable of shielding vulnerable epitopes [7]. IFV’s use of the latter strategy is seldom considered when predicting escape mutations despite the fact it has occurred dozens of times in the recorded history of human IFVs [25]. Given IFV’s propensity for shielding epitopes with glycans, predictive software capable of identifying sites likely to become glycosylated in the future may benefit those trying to develop the most effective models for predicting antigenic escape. Here we describe a simple albeit effective computational method for predicting future glycosylation sites on viral surface proteins. The method identifies which sites on viral proteins are most likely to mutate into N-linked glycosylation sequons based on the region’s nucleotide sequence, its surface accessibility, and the historical mutation frequency of residues within the region that would need to change to generate a glycosylation sequon. The method’s efficacy is then tested on historic influenza HA proteins and then applied to currently circulating strains.

## Materials and Methods

### Program overview

The method for predicting sequon emergence, *Future Sequon Finder* (FSF), was implemented in the Eclipse Java IDE under JDK 21 [26]. The general workflow of the program is outlined in Figure 1. It accepts a nucleotide sequence input with the option to include the associated surface accessibility for each AA in the protein acquired from the external surface accessibility prediction software ‘GetArea’ [27] as well as site-specific AA mutation frequencies acquired from the sequence alignment and analysis tool ‘MEGA’ using the Le and Gascuel model [28, 29]. The nucleotide input is converted into an AA sequence, cross-referencing the ‘GetArea’ results with the generated AA sequence, and a list of existing N-linked glycosylation sequons is identified along with lists of sites that are most likely to become glycosylation sequons as a result of genetic drift. Probable future glycosylation sites, or ‘near-sequons,’ are sorted into three categories based on the nucleic and amino acid edit distance from a glycosylation sequon and any changes in the physical properties of the AA sequence that would emerge from sequon transformation:

**Fig 1.**
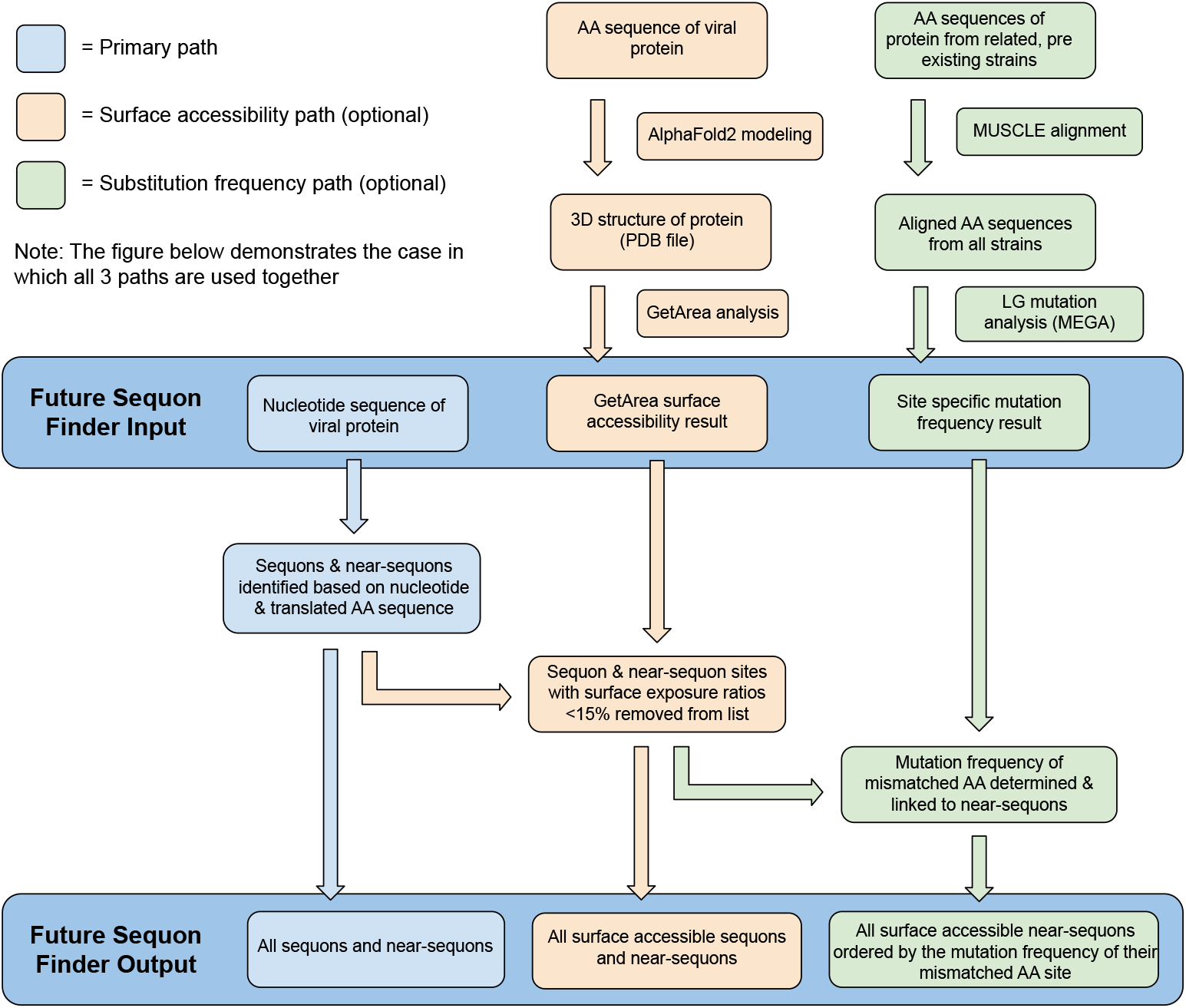
General workflow for generating outputs with Future Sequon Finder. The primary analytic pipeline shown in blue is the most basic analytic path FSF can follow, requiring only a nucleic acid sequence to generate lists of existing and near sequons. This diagram illustrates a case in which the mutation frequency (green) and surface accessibility (orange) data are combined with the primary path to generate more informative results. When using all pathways the FSF output will contain a a set of binned lists of: all near-sequons and true sequons, all near-sequons and true sequons with exposed surfaces, all near-sequons sorted by mutation frequency (not shown in the figure), and all near-sequons with solvent exposed surfaces sorted by mutation frequency.

- **Near-sequon** - A trio of amino acids in which only one AA must be changed to generate a glycosylation sequon (AA edit distance = 1)
- **Very near-sequon** - A trio of amino acids in which only one nucleotide must be changed to generate a glycosylation sequon (NT edit distance = 1)
- **Ultra near-sequon** - A very near-sequon wherein the mismatched AA has the same charge, polarity, and hydrophobicity as the correct AA

If surface accessibility data is included with the sequence input, all near-sequons will be sorted by the surface exposure ratio of the first AA in the sequon triad. All near-sequons (including ‘very’ and ‘ultra’ near-sequons) with exposure ratios greater than 15% are tagged as near-sequons with exposed surfaces, meaning they are adequately exposed for glycan attachment. While glycans are covalently bound to proteins prior to folding, we assume that a sequon in an unexposed region would either remain unglycosylated, thus providing no selective advantage, or become glycosylated and disrupt proper folding of the protein. If the user has chosen to incorporate site-specific mutation frequency data, near-sequons in each category with exposed surfaces will be sorted again based on the relative mutation frequency of the site where their mismatched AA is located. We assumed as a premise that if a near-sequon was only one AA away from becoming a true sequon, and that AA position was subject to frequent observable mutations in the history of related IFVs, that site was more likely to mutate again and generate a functional sequon. After each category is sorted, the rankings of all near-sequons are displayed to the user in their respective categories with those whose mismatched AA is most likely to mutate ranked at the top of the list.

### Sidechain exposure ratio analysis

To implement the surface accessibility path in FSF, the nucleotide sequence of interest was first be converted into AAs. A 3D protein model is then generated from the AA sequence and stored as a PDB file. ColabFold v1.5.5 (AlphaFold2 using MMseqs2) was used in this analysis to generate all 3D protein structures [30, 31]. After the PDB file is generated, the sidechain exposure ratios are determined. While there are many methods to achieve this, ‘GetArea’ was chosen due to its accessibility and ease of use [32, 33]. We used a 1.4 angstrom probe radius in GetArea for all predictions, the approximated radius of a water molecule. FSF was designed to receive GetArea outputs, and will assign a sidechain exposure ratio to each AA found in both the GetArea result and the converted nucleotide sequence input by the user. All AA/position identifier pairings from the GetArea result are compared to those same pairings generated from the converted nucleotide sequence input. If any mismatches between the GetArea sequence and user input sequence are detected, an error is thrown indicating the sidechain exposure ratios have not been properly aligned.

Once sidechain exposure ratios have been aligned to the converted AA sequence, FSF iterates through all categories of near-sequons previously determined from the nucleic acid sequence. The program sorts all near-sequons into their equivalent ‘near-sequon with exposed surface’ category if the sidechain exposure ratio of the first AA in the triad (asparagine, or the residue that would need to become asparagine) exceeds 15%. A cutoff of 15% was chosen given the observation that existing sequons seldom support glycans when the asparagine sidechain exposure ratios are less than 20% in HA proteins. This was determined using data from a study by Altman et al in which the glycosylation of sequons in influenza HAs was analyzed over time in conjunction with sidechain exposure results obtained using the pathway described above [25]. The additional 5% leniency was included to account for minor errors in 3D protein structure generation, and conformational changes that may occur during the transition from near-sequon to sequon.

### Site-specific mutation rate analysis

To implement the site-specific mutation frequency path in FSF, a list of nucleotide sequences relating to and predating the sequence of interest was first obtained. These sequences provide evolutionary context for the sequence of interest showing which regions are most likely to change over time. We excluded sequences with the following characteristics: uncertain nucleotides (any character other than A, T, G, or C), any AA insertions (HA sequences encoding more than 566 AAs), any repeat sequences sharing the same name, and any sequences whose collection year lay outside of a range specified by the user. The remaining nucleotide sequences were converted into AAs, and a FASTA file was generated and then aligned using the multiple sequence alignment tool MUSCLE [34]. The aligned sequences were saved as a .meg file and were imported into the Molecular Evolutionary Genetics Analysis (MEGA) tool [28]. Site-specific mutation rates were calculated in MEGA using the Le and Gascuel model with 8 gamma distribution categories [29].

The MEGA output was then imported into FSF, which assigned relevant site-specific mutation values to each predetermined near-sequon based on the AA position within that near-sequon that would need to change to generate a functional N-linked glycosylation sequon. For instance, if near-sequon ‘NTK’ was identified with AAs in positions 1, 2, and 3 respectively, FSF would recognize that the ‘K’ AA in position 3 must change to generate a sequon. FSF will then associate the site-specific mutation frequency for AA site 3 with that near-sequon. Near-sequons are then ranked within each classification category based on their associated mutation frequencies.

## Results

### Predicting historic IFV sequon addition events

To test the efficacy of the method implemented in FSF, a set of historic influenza strains with known SAEs in subsequent, closely related strains were obtained from Altman et al [25]. For this analysis, 20 strains of H1N1 viruses and 18 strains of H3N2 viruses relevant to SAEs were selected. Most of these sequences were selected from Figure S3 in a manuscript by Altman et al. who had already determined the phylogenetic relationships between prominent historic IFV strains related to SAEs [25]. Exceptions to this include A/NewYorkCity/2/1918, A/Tonga/14/1984, A/Memphis/1/1984, A/Colorado/14/2015, A/Virginia/22/2016, and A/Netherlands/10100/2024 for H1N1, and A/Wyoming/11/2014, A/Germany/13247/2022, A/Bangkok/ P3993/2023, A/Netherlands/10098/24 for H3N2 which were added to fill time gaps in the dataset and highlight important SAEs. All added strains were first selected from either the Bacterial and Viral Bioinformatics Resource Center (BV-BRC) [35] or GISAID’s EpiFlu database [36] before being compared to other circulating strains from their time to ensure the selected strains were representative of common glycosylation sequon (and near-sequon) patterns rather than rare variants. The strains, sequences, and their source can be found in Supplemental Material.

The nucleic acid sequences for the HA proteins of all 38 selected strains were obtained from the BV-BRC [37] and GISAID’s EpiFlu database [38]. The AA-specific sidechain exposure ratios for each IFV strain were determined using AlphaFold2 and GetArea as above. HAs were analyzed by GetArea in their monomeric forms to obtain sidechain exposure, and therefore the number of unexposed sites is likely an underestimate compared to the number found in trimeric HAs. To determine site-specific mutation frequencies, HA nucleotide sequences were obtained from BVBRC and GISAID (32,000 H1N1 sequences, 40,000 H3N2 sequences) to form H1N1 and H3N2 master lists. The mutation frequencies were calculated separately for the analysis of every strain preceding an SAE using only strains that predated or coincided with the analysis strain. For example, site-specific mutation frequencies for A/Udron/307/1972(H3N2) were calculated only using strains from the H3N2 master set that circulated prior to 1972. Due to computational restrictions, if a single year had more than 500 associated strains, 500 were randomly selected from that year to contribute to the mutation frequency analysis. For years with strain numbers below the cutoff, all strains were used. The number of IFV HA genomes obtained for each year is shown in Figure 2.

**Fig 2.**
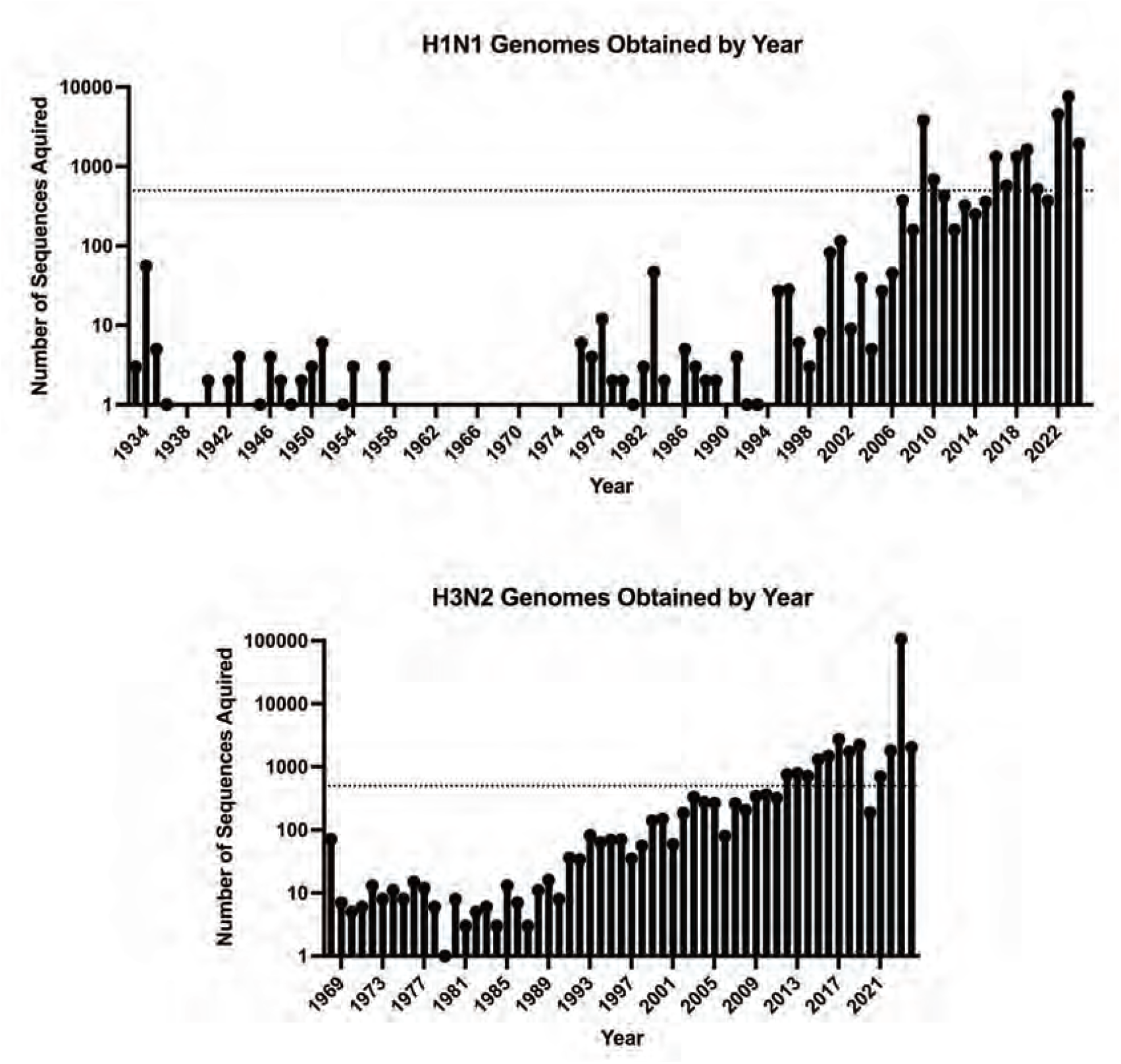
Influena genomes obtained by year. These histograms represent the total number of sequences in the H1N1 and H3N2 master lists acquired from the BV-BRC [37] and GISAID’s EpiFlu database [38] for each year. The dotted line at Y=500 represents the cutoff for mutation frequency analysis that was imposed due to computational limits.

For each strain that would be analyzed, the nucleotide sequence was uploaded into FSF along with its associated sidechain exposure and mutation frequency data. In this study, we focused on sequon alterations in the H1 head domain of the HA given that this region is where most changes in N-linked glycosylation occur. We ignored sequons and near-sequons outside this region to decrease the total number of near-sequons in the outputs and computational burden. Full-length HAs were used to generate all sidechain exposure and mutation frequency data.

We also analyzed swine H1N1 strains. Six thousand swine H1N1 sequences collected from 1931-2022 were obtained from the BV-BRC [35] and pre-processed using a custom Java program designed to identify SAE strains. Once identified, a maximum-likelihood phylogenetic tree containing 15 SAE strains and 700 other H1N1 swine strains (a maximum of 25 strains selected from each year, 1931-2022) was generated and close relatives to the SAE strains were identified. Those close relatives predating the SAE were analyzed with FSF with their accompanying sidechain exposure and mutation frequency data as described for the human influenza strains. Unlike the analysis of human IFVs, rare sequons were not excluded from the swine analysis. The only swine SAE strains to be excluded from the analysis were those with no close ancestors defined as those strains on isolated branches of the phylogenetic tree.

### Predicting future IFV sequon addition events

Ten currently circulating IFV strains were selected for both H1N1 and H3N2 from GISAID’s EpiFlu database. These strains were selected semi-randomly, with each strain obtained from a different nation to cover a wider range of circulating influenza geographic diversity. The selected strains include:

- **H1N1** - A/Netherlands/10114/2024, A/Curacao/10079/2024, A/Aragon/102/2024, A/Bayern/33/2024, A/SriLanka/32/2024, A/Bangkok/P479/2024, A/Timis/563634 /2024, A/Norway/01013/2024, A/NorthCarolina/14611/2024, A/UnitedKingdom/ 14762/2024
- **H3N2** - A/Curacao/10082/2024, A/Canberra/14/2024, A/Bangkok/P458/2024, A/ Hungary/1/2024, A/Netherlands/10110/2024, A/Norway/00732/2024, A/Victoria /12/2024, A/Philippines/2/2024, A/Yekaterinburg/2/2024, A/Oklahoma/14622/2024

Each chosen strain was analyzed using all FSF pathways. Sidechain exposure ratios were calculated for each strain individually and mutation frequencies were calculated with the same HA sequence dataset and ‘pick 500’ approach used for the historic strain analysis. For H1N1, mutation frequencies were calculated using strains from all years (1918-2024). Mutation frequencies were re-calculated with strains from 2009-2024 to determine if there was a noteworthy difference between the resultant ranking of near-sequons given the large phylogenetic distance between modern H1N1 strains and those circulating pre-2009. For H3N2 viruses the mutation frequency was calculated once using strains from all years (1968-2024). All exposed very near-sequons found in the circulating IFV strains were assigned an overall rank based on the relative mutation rate of their mismatched AA site. The frequency of the very near-sequon sites in the analyzed IFVs was reported but not factored into the overall ranking.

Initially, FSF was designed to only consider the genetic composition of HA sequences. Following the primary program path in Figure 1, FSF was able analyze the selected 20 H1N1 and 18 H3N2 strains with their nucleotide sequences. Sites that would become glycosylation sequons at some point in the IFV timeline were tracked over time. Their status as existing sequons, near-sequons, very near-sequons, and ultra near-sequons was recorded (Table 1).

**Table 1.**
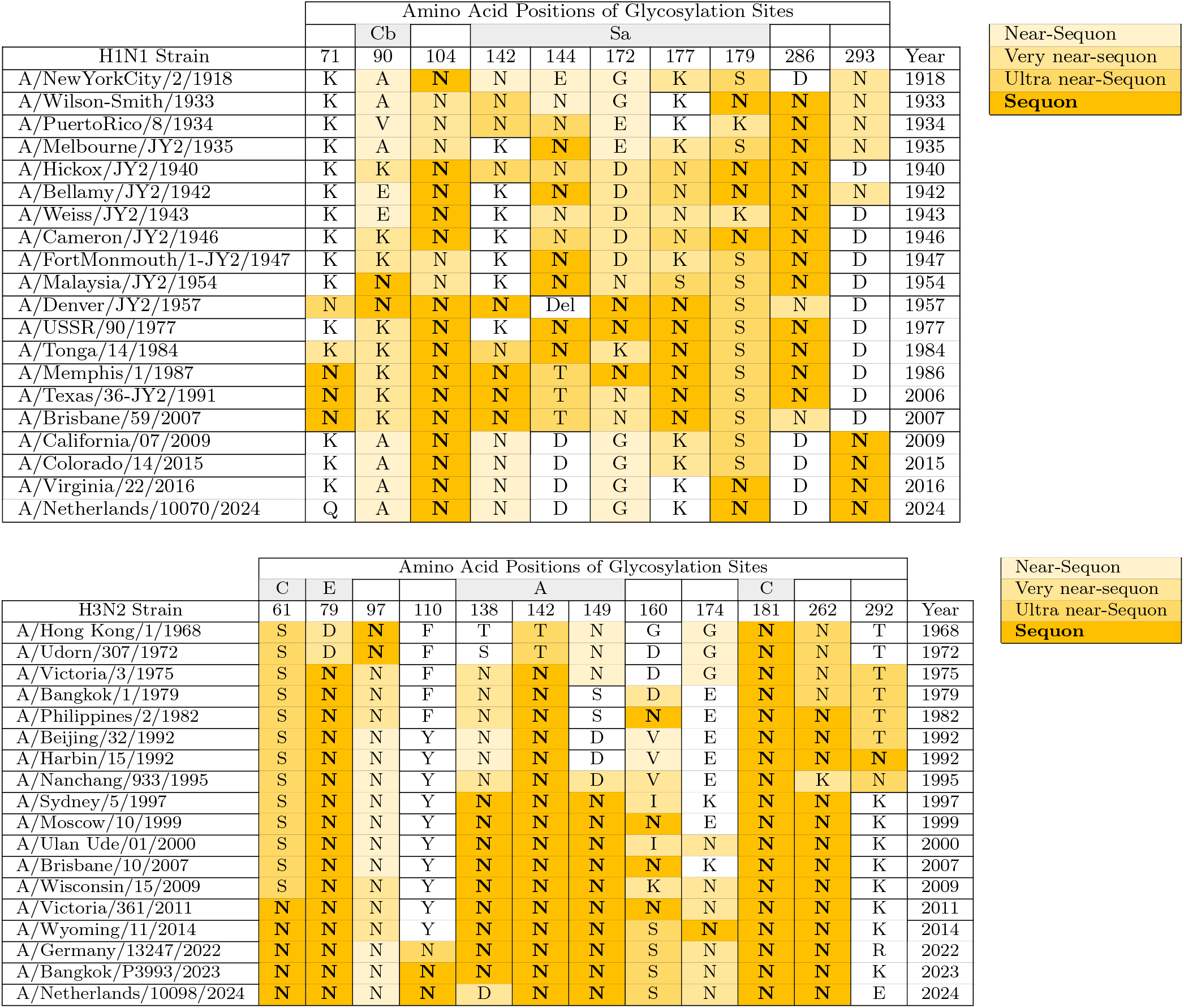
Temporally organized FSF results in the context of historically glycosylated HA regions. Historic H1N1 and H3N2 IFV strains have been temporally organized to illustrate the trends in the glycosylation patterns over time as well as the trends in near-sequon occurrence in these critical sites. The letters in each AA site column are the single-letter identifiers for the AA found at the given position in the given strain. These single-letter identifiers also represent the first AA of the sequon (or near-sequon) triad (the ‘N’ position of N-!P-S/T). Characterized antigenic sites are indicated in the top row of the table. Near-sequons identified by FSF are indicated by varying shades of orange, with darker shades representing sites that more closely resemble glycosylation sequons. The emboldened letters in the darkest shade of orange represent true N-linked glycosylation sequons. (**Top**) Temporally organized H1N1 IFVs. (**Bottom**) Temporally organized H3N2 IFVs.

Considering the potential genetic disparity between IFV strains from consecutive years, a phylogenetic analysis of the same IFV HA sequences was conducted. This strategy avoids pairing distantly related sequences and captures the impact of genetic drift, the primary means by which IFVs adapt to escape herd immunity [7, 39], on sequon emergence. Phylogenetic trees containing all selected strains for H1N1 and H3N2 were constructed and are displayed in Figure 3. These trees were used to determine the branch on which each SAE occurred. Assuming parsimony, there were 19 SAEs in the H1N1 lineage and 13 SAEs in the H3N2 lineage that ultimately rose to prominence, persisting for *≥*2 years and successfully spreading to multiple countries. It is worth noting other SAEs have occurred in recorded human H1N1 and H3N2 HA sequences, however, these glycosylation sequons would appear in single countries and would disappear within 1-2 years. This analysis only considers SAEs that rose to prominence as previously defined. Preexisting strains closely related to the SAE strain were selected and analyzed with FSF. Table 2 displays the results of this phylogenetic-based analysis. To better visualize the disparity between HA sequences within IFV groups over time, AA edit distance heatmaps were generated using randomly selected HA strains from the H1N1 and H3N2 master lists (Figure 4).

**Table 2.** FSF site designations of critical sites from closely related IFV strains circulating before SAEs. All AA sites listed in this table become N-linked glycosylation sequons in strains listed in the ‘SAE Strain’ column. Phylogenetically related IFV strains circulating before the SAE have been selected and the FSF designation of the site that would become glycosylated is stated in the rightmost column. Related strains were selected if they predated the strain in which the glycosylation sequon arose and was the closest preexisting relative on the generated phylogenetic tree shown in Figure 3.

| Group | Strain name | Site of SAE | Strain circulating prior to SAE | Near-sequon Designation |
| --- | --- | --- | --- | --- |
| H1N1 | A/WilsonSmith/1993 | 179 | A/NewYorkCity/2/1918 | Ultra near-sequon |
|  | A/WilsonSmith/1993 | 286 | A/NewYorkCity/2/1918 | Not detected |
|  | A/Melbourne/JY2/1935 | 144 | A/PuertoRico/8/1934 | Ultra near-sequon |
|  | A/Bellamy/JY2/1942 | 104 | A/Melbourne/JY2/1935 | Very near-sequon |
|  | A/Bellamy/JY2/1942 | 179 | A/Melbourne/JY2/1935 | Ultra near-sequon |
|  | A/Hickox/JY2/1940 | 179 | A/Melbourne/JY2/1935 | Ultra near-sequon |
|  | A/FortMonmouth/1-JY2/1947 | 144 | A/Cameron/JY2/1946 | Ultra near-sequon |
|  | A/Malaysia/JY2/1954 | 90 | A/FortMonmouth/1-JY2/1947 | Very near-sequon |
|  | A/Denver/JY2/1957 | 104 | A/Malaysia/JY2/1954 | Very near-sequon |
|  | A/Denver/JY2/1957 | 142 | A/Malaysia/JY2/1954 | Not detected |
|  | A/Denver/JY2/1957 | 172 | A/Malaysia/JY2/1954 | Very near-sequon |
|  | A/Denver/JY2/1957 | 177 | A/Malaysia/JY2/1954 | Ultra near-sequon |
|  | A/USSR/90/1977 | 144 | A/Cameron/JY2/1946 | Ultra near-sequon |
|  | A/USSR/90/1977 | 172 | A/Cameron/JY2/1946 | Very near-sequon |
|  | A/USSR/90/1977 | 177 | A/Cameron/JY2/1946 | Ultra near-sequon |
|  | A/Memphis/1/1987 | 71 | A/Tonga/14/1984 | Very near-sequon |
|  | A/Memphis/1/1987 | 142 | A/Tonga/14/1984 | Ultra near-sequon |
|  | A/California/07/2009 | 293 | A/NewYorkCity/2/1918 | Very near-sequon |
|  | A/Virginia/22/2016 | 179 | A/Colorado/14/2015 | Ultra near-sequon |
| Group | Strain Name | Site of SAE | Strain circulating prior to SAE | Near-sequon Designation |
| H3N2 | A/Victoria/3/1975 | 79 | A/Udorn/307/1972 | Very near-sequon |
|  | A/Victoria/3/1975 | 142 | A/Udorn/307/1972 | Ultra near-sequon |
|  | A/Philippines/2/1982 | 160 | A/Bangkok/1/1972 | Very near-sequon |
|  | A/Philippines/2/1982 | 262 | A/Bangkok/1/1972 | Ultra near-sequon |
|  | A/Harbin/15/1992 | 292 | A/Beijing/32/1992 | Ultra near-sequon |
|  | A/Sydney/5/1997 | 138 | A/Nanchang/933/1995 | Very near-sequon |
|  | A/Sydney/5/1997 | 149 | A/Nanchang/933/1995 | Very near-sequon |
|  | A/Sydney/5/1997 | 262 | A/Nanchang/933/1995 | Very near-sequon |
|  | A/Moscow/10/1999 | 160 | A/Sydney/5/1997 | Very near-sequon |
|  | A/Victoria/361/2011 | 61 | A/Wisconsin/15/2009 | Ultra near-sequon |
|  | A/Victoria/361/2011 | 160 | A/Wisconsin/15/2009 | Very near-sequon |

**Fig 3.**
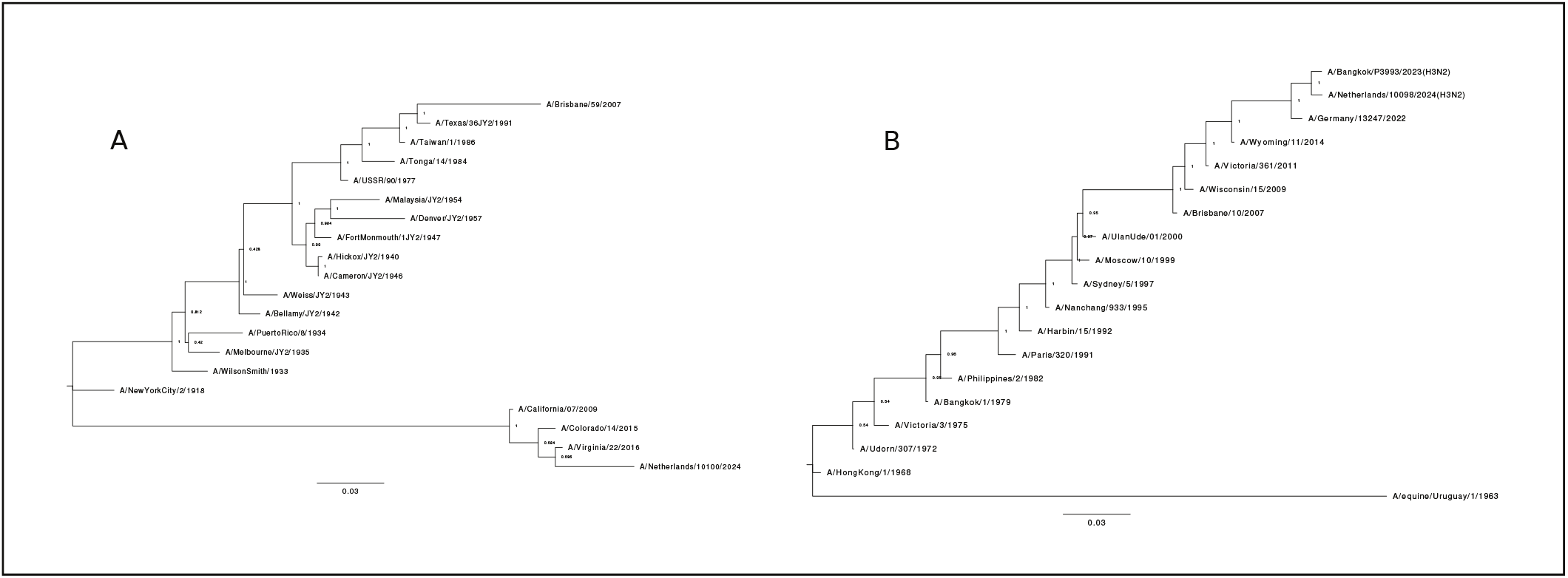
Phylogenetic trees containing selected H1N1 and H3N2 IFV HAs. HA nucleotide sequences for each of the 20 H1N1 and 17 H3N2 IFVs chosen for historical analysis were aligned and used to generate a maximum likelihood phylogenetic tree in MEGA [28]. Bootstrapping (500x) values are displayed at each branch node as a proportion. (**A**) Maximum likelihood tree of selected H1N1 IFVs. (**B**) Maximum likelihood tree of selected H3N2 IFVs. A/equine/Uruguay/1/1963 was used as an outgroup to root this tree.

**Fig 4.**
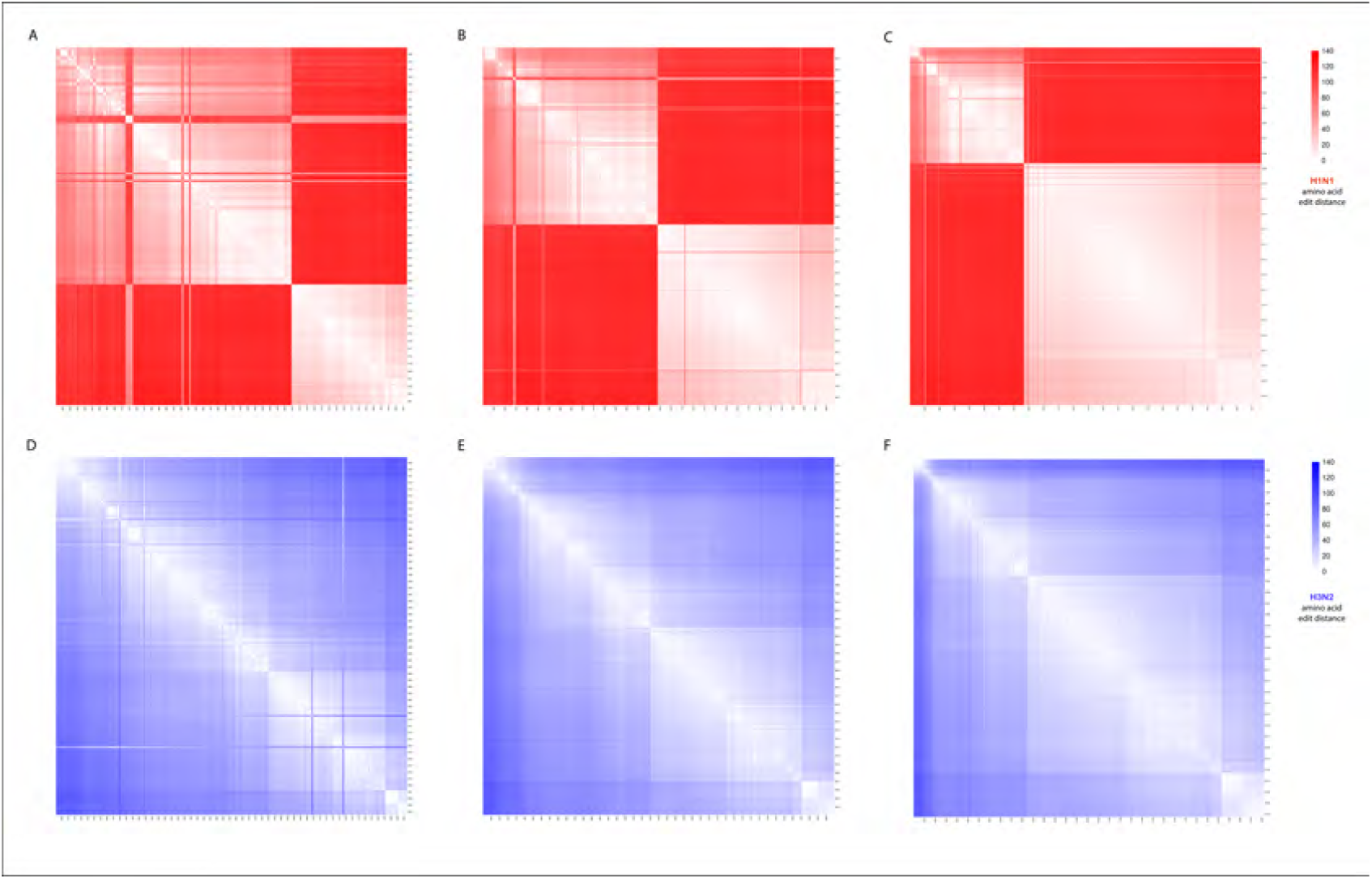
Mapping edit distance between IFV HAs over time. To better understand the roles of antigenic shift and antigenic drift in H1N1 and H3N2 HA evolution, edit distances between HA AA sequences were calculated and visualized in R studio. All HA sequences were randomly selected from the master lists comprised of 32,000 H1N1 sequences and 40,000 H3N2 sequences. As the number of sequences drawn per year increases, modern strains represent a larger portion of the heat map given the increased number of sequences available. (**A**) H1N1 HA protein AA edit distances from 1918-2024, maximum of 5 strains per year. (**B**) H1N1 HA protein AA edit distances, maximum of 25 strains per year. (**C**) H1N1 HA protein AA edit distances, maximum of 100 strains per year. (**D**) H3N2 HA protein AA edit distances from 1968-2024, maximum of 5 strains per year. (**E**) H3N2 HA protein AA edit distances, maximum of 25 strains per year. (**F**) H3N2 HA protein AA edit distances, maximum of 100 strains per year.

While FSF had proven to be effective at tagging sites as near-sequons in strains circulating before SAEs, it would also tag many sites that would never become sequons. To eliminate some of these false positives, the side chain surface exposure analysis method was implemented. The site-specific mutation frequency analysis method was implemented to further organize the remaining near-sequons with exposed sidechains. At this stage, FSF was using all three pathways outlined in Figure 1 to generate the historic results displayed in Table 3. Strain comparisons matched those used in the phylogenetic analysis.

**Table 3.**
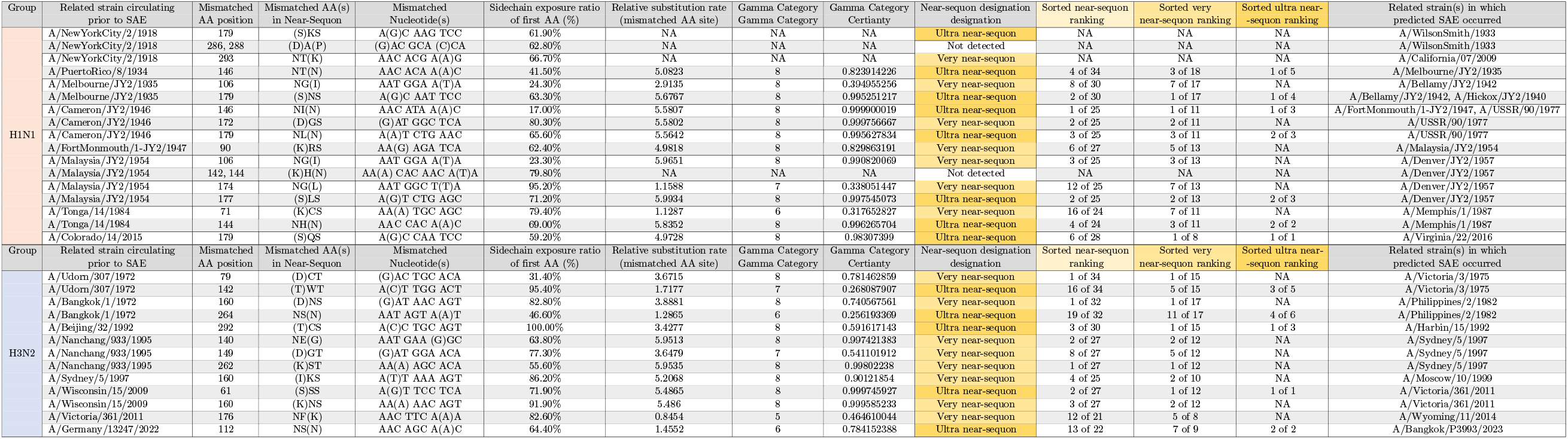
Historic FSF predictions with side-chain exposure and site-specific mutation frequency data. The same closely related H1N1 and H3N2 IFV strains selected for analysis in Figure 2 are analyzed again in conjunction with sidechain exposure and site-specific mutation frequency data. The surface exposure of the first AA in each near sequon is expressed as a percentage taken from the GetArea result. The relative mutation rate of the mismatched AA position is displayed, with a value of 1 representing the average mutation rate for AAs across the HA protein calculated for that period. The site-specific mutation gamma category placement out of 8 is displayed, with AAs assigned to category 8 being the most likely to change. The certainty with which MEGA assigned sites to their given gamma categories is also displayed as a proportion. The rankings of near-sequons among others in the same HA are displayed based on relative AA mutation rates. Near-sequons with surface exposure ratios *≤*15% are excluded from the rankings.

Curious to see how far the predictive capacity of FSF extended, we tested its ability to predict the emergence of glycosylation sequons in swine H1N1 strains as well. This analysis focuses on all swine H1N1 HA SAEs from 1931-2022 where the SAE strain has a close ancestor suitable for FSF analysis recorded in the BV-BRC [35]. The results of that analysis are displayed in Table 5.

Following the predictive success FSF displayed with the historic IFV datasets, it was time to generate future predictions from currently circulating strains. All accurate SAE predictions in the historic human dataset were predicted as very near-sequons or ultra near-sequons, so it is logical to feature these categories when predicting future SAEs. Predicted sites were selected and sorted by FSF using both surface accessibility and mutation frequency data. Final rankings were determined by site-specific mutation frequency alone. The summarized results of these future sequon predictions are displayed in Table 4.

**Table 4.** Future H1N1 and H3N2 N-linked glycosylation sequon locations predicted by FSF. Each table summarizes the top site predictions from 10 individual FSF runs with currently circulating IFV HAs. Overall near-sequon rankings were determined by their site-specific mutation rates. Near-sequon frequencies are displayed as proportions, along with the average sidechain exposure ratios of the first Aas in the near-sequon triads. The mutation rates for the mismatched AA positions along with their gamma category assignments are displayed similarly (**Top**) Future H1N1 sequon locations predicted by FSF using a mutation frequency calculation based on a random selection of *≤* 500 H1N1 HA sequences per year for all years from 1918-2024. (**Middle**) Future H1N1 sequon locations predicted by FSF using a mutation frequency calculation based on a random selection of *≤* 500 H1N1 HA sequences per year for all years from 2009-2024. (**Bottom**) Future H3N2 sequon locations predicted by FSF using a mutation frequency calculation based on a random selection of *≤* 500 H3N2 HA sequences per year for all years from 1968-2024.

| H1N1 All Years |  |  |  |  |  |  |  |  |  |  |
| --- | --- | --- | --- | --- | --- | --- | --- | --- | --- | --- |
| Overall ranking | Near sequon site | Near-sequon frequency | Average sidechain exposure | Mutation rate | Gamma category | Gamma certainty | Near-sequon designation | Average NS ranking | Average VNS ranking | Average UNS ranking |
| 1 | 286 | 0.1 | 64.50% | 4.6243 | 8 | 0.999991929 | Very near-sequon | 7 | 1 | NA |
| 2 | 111 | 0.1 | 85.10% | 4.5318 | 8 | 0.961737947 | Very near-sequon | 8 | 1 | NA |
| 3 | 111 | 0.1 | 78.60% | 1.3607 | 6 | 0.744258253 | Very near-sequon | 11 | 1 | NA |
| 4 | 201 | 1 | 51.18% | 1.3296 | 6 | 0.711083036 | Ultra near-sequon | 11.6 | 1.3 | 1 |
| 5 | 136 | 1 | 31.66% | 1.0194 | 5 | 0.510145247 | Very near-sequon | 12.6 | 2.3 | NA |
| 6 | 245 | 1 | 21.52% | 0.9383 | 5 | 0.465989051 | Very near-sequon | 14.6 | 3.3 | NA |
| 7 | 256 | 1 | 65.71% | 0.6547 | 4 | 0.471340697 | Very near-sequon | 15.6 | 4.3 | NA |
| 8 | 181 | 0.1 | 37.70% | 0.5096 | 4 | 0.544519551 | Ultra near-sequon | 18 | 6 | 2 |
| 9 | 73 | 1 | 26.31% | 0.25 | 3 | 0.42380528 | Very near-sequon | 21.5 | 5.4 | NA |

| H1N1 2009+ |  |  |  |  |  |  |  |  |  |  |
| --- | --- | --- | --- | --- | --- | --- | --- | --- | --- | --- |
| Overall ranking | Near sequon site | Near-sequon frequency | Average sidechain exposure | Mutation rate | Gamma category | Gamma certainty | Near-sequon designation | Average NS ranking | Average VNS ranking | Average UNS ranking |
| 1 | 286 | 0.1 | 64.50% | 4.3548 | 8 | 0.999946882 | Very near-sequon | 4 | 1 | NA |
| 2 | 111 | 0.1 | 85.10% | 2.3362 | 7 | 0.832885697 | Very near-sequon | 10 | 1 | NA |
| 3 | 136 | 1 | 31.90% | 1.5147 | 6 | 0.526586082 | Very near-sequon | 11.2 | 1.2 | NA |
| 4 | 201 | 1 | 51.14% | 1.3234 | 6 | 0.648165763 | Ultra near-sequon | 12.2 | 2.2 | 1 |
| 5 | 256 | 1 | 65.72% | 1.0935 | 6 | 0.494117309 | Very near-sequon | 14.2 | 3.2 | NA |
| 6 | 111 | 0.1 | 78.60% | 1.0706 | 6 | 0.526596067 | Very near-sequon | 15 | 4 | NA |
| 7 | 181 | 0.1 | 37.70% | 0.6317 | 4 | 0.434216146 | Ultra near-sequon | 16 | 5 | 2 |
| 8 | 245 | 1 | 21.47% | 0.6194 | 4 | 0.362708452 | Very near-sequon | 15.7 | 4.4 | NA |
| 9 | 73 | 1 | 26.38% | 0.1835 | 2 | 0.422196462 | Very near-sequon | 21.5 | 5.4 | NA |

| H3N2 All Years |  |  |  |  |  |  |  |  |  |  |
| --- | --- | --- | --- | --- | --- | --- | --- | --- | --- | --- |
| Overall ranking | Near sequon site | Near-sequon frequency | Average sidechain exposure | Mutation rate | Gamma category | Gamma certainty | Near-sequon designation | Average NS ranking | Average VNS ranking | Average UNS ranking |
| 1 | 160 | 1 | 84.43% | 5.2486 | 8 | 0.999979796 | Ultra near-sequon | 2 | 1 | 1 |
| 2 | 138 | 0.5 | 67.26% | 5.1587 | 8 | 0.969068272 | Very near-sequon | 4 | 2 | NA |
| 3 | 174 | 1 | 79.49% | 3.4833 | 7 | 0.607051602 | Very near-sequon | 5.5 | 2.5 | NA |
| 4 | 223 | 0.9 | 63.66% | 2.3371 | 7 | 0.992646267 | Very near-sequon | 7.6 | 3.6 | NA |
| 5 | 206 | 1 | 44.09% | 2.3287 | 7 | 0.987549431 | Very near-sequon | 8.5 | 4.4 | NA |
| 6 | 213 | 0.1 | 44.10% | 2.2362 | 7 | 0.891834972 | Very near-sequon | 10 | 5 | NA |
| 7 | 280 | 1 | 94.70% | 2.1309 | 7 | 0.783493861 | Very near-sequon | 11.5 | 5.5 | NA |
| 8 | 121 | 1 | 40.17% | 1.4466 | 6 | 0.679889132 | Very near-sequon | 12.5 | 6.5 | NA |
| 9 | 140 | 0.2 | 100.00% | 1.2228 | 6 | 0.693254615 | Ultra near-sequon | 13 | 6.5 | 2 |
| 10 | 69 | 0.9 | 42.13% | 0.7351 | 5 | 0.381898704 | Very near-sequon | 15.7 | 7.8 | NA |
| 11 | 175 | 1 | 88.49% | 0.4438 | 4 | 0.452966492 | Very near-sequon | 18.5 | 8.6 | NA |
| 12 | 168 | 0.6 | 16.97% | 0.0618 | 1 | 0.573958363 | Very near-sequon | 20.8 | 9.7 | NA |

**Table 5.**
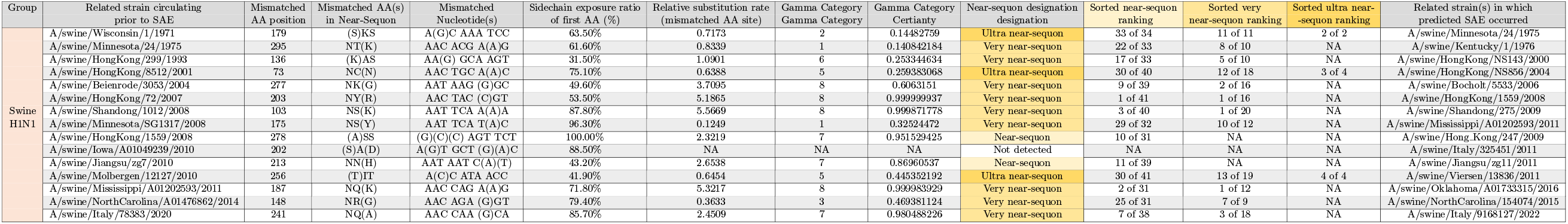
Historic Swine H1N1 FSF predictions. This table shows FSF results from swine IVFs closely related to SAE strains with sidechain exposure and site-specific mutation frequency data. The layout of this table is identical to Table 3. As was the case in the analysis of human IFVs, near-sequons with surface exposure ratios *<*15% are excluded from the rankings.

## Discussion

The temporal analysis of historic H3N2 IFVs supported the notion that the HA proteins of currently circulating strains are often most closely related to the HAs of strains circulating in the previous year (Figures 3 and 4). The temporal analysis with H1N1 revealed a similar pattern with some exceptions, given that strains closely related to bygone IFVs occasionally reemerged and usurped the predominant strains. The most dramatic example was the emergence of A(H1N1)pdm09 type IFVs from swine reservoirs in 2009, whose HA proteins and glycosylation patterns were more closely related to those from the 1976 swine outbreak than any circulating in the prior decade. This temporal analysis of amino acid and glycosylation changes in H1N1 strains demonstrates the importance of monitoring a variety of IFVs that may be surviving at low levels in human populations, or even propagating in animal reservoirs waiting for an opportunity re-emerge. Our results suggest, however, that for both H1N1 and H3N2 that prominent circulating strains often serve as the best FSF inputs for predicting sequons that will arise in the near future.

Through the phylogenetic analysis of historic IFV strains with FSF, we found that that N-linked glycosylation sites on HA proteins almost always arise due to single nucleotide mutations. Of the observed 30 SAEs in the historic IFV selection, 28 arose due to single nucleotide mutations. Of the two that did not, one appeared to be the result of a deletion that occurred in A/Denver/JY2/1957 (compared to A/Malaysia/JY2/1954) at AA position 144, followed by two single nucleotide changes, one in AA 142 and the other in AA 145 that would take position 144 following the deletion event. The other appeared to be due to two single nucleotide mutations in A/WilsonSmith/1933 (compared to A/NewYorkCity/2/1918), one in AA codon 286 and the other in AA 288. In both cases, these mutations may have arisen simultaneously to generate glycosylation sequons. Alternatively, it could be that more closely related strains containing near-sequons at these sites were never sequenced given the limited number of HA sequences available from the early 20th century. The notion that N-linked glycosylation sequons almost always arise due to single nucleotide mutations coincides with the finding that antigenic diversity in IFV HAs is primarily driven by the gradual accumulation of single base pair mutations [40, 41]. Though unsurprising, this finding allows for more accurate sequon predictions given that sites requiring more than a single base mutation can be eliminated from consideration. While it is clear that the ultra and very near-sequon categories have a significantly different proportion of SAEs than the near-sequon category, 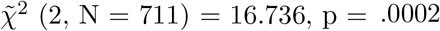, the difference between the ultra and very near-sequon categories is less apparent, 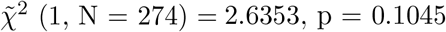. More data is needed to determine if this difference is statistically significant.

Our analysis also noted a gap in the H1N1 HA sequence coverage from 1957-1976 shown in Figure 2. This gap is explained by the emergence of H2N2 viruses in 1957 which quickly became the dominant circulating IFVs until disappearing a decade later, replaced by H3N2 [42]. H1N1 viruses would remain in the shadows until a swine variant would emerge and cause a pandemic in 1976, followed by historic H1N1 strains returning to prominence in 1977. Fascinatingly, the H1N1 strains circulating in 1977 were very closely related to those circulating in the 1950s, so much so that it appeared as if almost no genetic drift had occurred over the decades between the strains [43, 44]. This helps to explain why FSF was still able to predict the SAEs in sites 144, 172, and 177 that occurred in the A/USSR/90/1977 H1N1 strain when analyzing A/Cameron/JY2/1946 despite the large time gap between the sequences. In actuality, these glycosylation sequons arose in the 1950s in strains like A/FortLeonardWood/1951 and the same glycosylation patterns likely reemerged in the 1970s through the reemergence of these historic strains themselves [43, 44]. The H1N1 HA edit distance heatmaps in Figure 4 show this history, including the emergence of a novel swine-origin H1N1 strain in 1976 closely resembling modern H1N1pdm09 HAs followed by the emergence of strains closely resembling those predating the 1976 pandemic. H1N1pdm09-like Has arose again in several strains such as A/Ohio/3559/1988 and A/Maryland/12/1991 before becoming the dominant H1N1 HA types after the 2009 pandemic. Juxtaposed to the H1N1 heatmaps, the evolution of H3N2 HAs is gradual with the most noticeable shifts occurring in 2003 and 2021.

Following the phylogenetic analysis, the reduction and sorting of near-sequon pools became a priority leading to the implementation of the two optional pathways shown in figure 1. The sidechain exposure ratio analysis of individual AA sites greatly narrowed the near-sequon pools. A typical implementation of the surface accessibility pathway reduces HA near-sequon pools by 35-45% (^∼^50 near-sequons reduced to ^∼^30). In the analysis of historic human and swine IFVs, no near-sequon that would subsequently become a sequon was excluded by the 15% sidechain exposure cutoff, though some were close. To be safe in future analysis, it may be beneficial to slightly lower this cutoff. To explain the lack of N-linked glycosylation in unexposed regions, we propose that SAEs with unexposed asparagine residues either remain unglycosylated or become glycosylated before protein folding, introducing detrimental conformational changes. Given the co-translational nature of glycan addition via oligosaccharyltransferase, a region that is unexposed post-folding may be exposed at the time of glycosylation. Adding a bulky oligosaccharide moiety to such a site, however, would likely result in significant conformational changes during folding, rendering the HA protein ineffective. If the SAE site folds correctly into its unexposed state before glycan binding, glycosylation will not occur. As a result, the mutation will provide no selective advantage via epitope shielding and is less likely to rise to prominence.

In conjunction with the solvent accessibility results, the mutation frequency analysis proved a powerful predictor of which near-sequons would become true sequons in the human IFV dataset. In particular, ranking very near-sequons based on the mutation frequencies of their mismatched AAs resulted in almost all correct near-sequons being found at the top of their respective lists. Of the 25 correctly predicted very near-sequons, 24 were ranked in the top 10 very near-sequons, 19 in the top 5, and an impressive 17 were ranked in the top 3 based on site specific mutation frequency. It is also worth noting that all correctly predicted near-sequons with one exception had associated site-specific mutation rates exceeding 1.0, meaning the rates of the sites that would ultimately mutate and generate sequons were almost always greater than the average AA mutation rate across the HA. Interestingly, the mutation frequency analysis proved to be effective even when calculated from a minimal number of preexisting sequences, especially in the case of H1N1 viruses in the early to mid-20th century. H1N1 mutation analyses for 1934 and 1935 successfully placed all correct very near-sequons in the top 7, despite the calculation being based on only 59 and 64 preexisting H1N1 strains respectively. It is unclear if the abundance of available strains from the early 2000s and onwards significantly improves the power of mutation frequency analysis as a predictive tool, as there have been few SAEs in H1N1 and H3N2 since the turn of the century. While it seems reasonable that larger sequence datasets will improve the predictive power of mutation analyses, more data is needed.

In the case of the swine influenza dataset, near-sequon designations, along with sidechain exposure ratios, were effective predictive tools. Fourteen of 15 SAEs were tagged as near-sequons with exposed surfaces, with 12 of 14 being very near-sequons. Site-specific mutation frequency proved less effective in this dataset, with 50% of very near-sequons being ranked in the bottom half of the very near-sequon pool, compared to 15% in the human H1N1 dataset and 23% in the human H3N2 dataset. There are many differences between IFV circulation in swine and human populations, one of the most apparent being the transmission dynamics. Swine herds are often tightly regulated and have limited exposure to other herds, contrasted to humans who frequently encounter other individuals worldwide [45]. This may explain why SAEs in the swine dataset rarely rise to prominence, while SAEs in the human dataset often do. The isolation of swine herds may also lead to an increased emphasis on local IFV adaptation. This would imply mutation frequencies in one population may not align with those in other pockets of swine IFV adapting to their local environment, potentially explaining why the mutation frequency parameter was less effective in the swine dataset.

The historical IFV analysis successfully demonstrated the efficacy of FSF, but it also revealed weaknesses. FSF excels at predicting SAE locations when the analyzed strain is closely related to the SAE strain, and the SAE occurs in a region with a high mutation rate. It isn’t guaranteed that next year’s predominant IFVs will be most closely related to this year’s, though, particularly in the case of H1N1. In the event of historic strain reemergence, FSF analyses based on circulating strains distantly related to the emerging strain will be less accurate. This is also true in the event of novel strain emergence from animal reservoirs, as seen in the H1N1 strain from 2009. Furthermore, SAEs occasionally occur in regions with low mutation rates as was the case for the H3N2 SAE at position 176 in A/Wyoming/11/2014, with a site-specific mutation rate of only 0.85. Fortunately, these difficult-to-predict events rarely occur in the historical human dataset. It is also worth noting that FSF only predicts the emergence of sequons, and does not guarantee that those sequons will become glycosylated. The sidechain exposure analysis used by FSF may help to provide some insight into this, as unexposed sequons will seldom be glycosylated. There are, however, other factors influencing whether a glycan will bind to a sequon such as which amino acids surround the tripeptide sequon [46]. Fortunately, there are numerous existing programs designed to determine which sequons in a protein will become glycosylated [47–51]. Running the most probable near-sequon mutants determined by FSF through programs such as these may further enhance our ability to predict future glycosylation sites.

The analysis of currently circulating H1N1 and H3N2 IFV strains revealed sites that may become glycosylated in the future assuming future SAE strains are most closely related to those circulating presently. In the historical analysis of human IFVs, every correct near-sequon prediction was tagged as a very near-sequon. Consequently, only very-near sequons (or ultra near-sequons) were considered serious candidates for SAEs in the near future. All near-sequons appearing in the lists were the highest-ranking sites predicted by FSF. It is important to track these near-sequons over time as a near-sequon found in strains circulating this year may not be found in circulating strains next year, and vice versa. For example, position 142 in the H1N1 HA is not included in the list of predicted sites given that it is not a single nucleotide change away from becoming a sequon, but the mismatched AA in this near-sequon (position 144) currently has the highest mutation rate in the entire H1 domain of the H1N1 HA protein and may become a very near-sequon in time. For this reason, an analysis of presently circulating IFVs should be conducted every few years to catch the new very near-sequons that emerge. That being said, should a new glycosylation sequon emerge in the near future in either H1N1 or H3N2, it is likely included in the tables displayed in Figure 4.

The historic analysis of IFVs with FSF demonstrated that it is often possible to predict N-linked glycosylation sequon locations before they emerge in H1N1 and H3N2 HA proteins. Facinatingly, the predictive capacity seems to extend to swine H1N1 viruses as well though the mutation frequency parameter was less effective. Whether this predictive capacity extends to HA proteins in other IFV types, such as influenza B viruses or those circulating in avian reservoirs, should be tested. Glycosylation changes on the IFV neuraminidase surface protein are also likely to be predictable. This predictive capacity may even extend to other viral surface proteins such as the heavily glycosylated envelope glycoprotein of the human immunodeficiency virus (HIV) which is known to alter glycosylation patterns to take advantage of glycan shielding [52], or the glycosylated SARS-CoV-2 spike protein. FSF serves as proof of concept, and the predictive power of sequon prediction programs like it can undoubtedly be improved. Unfortunately, the list of historic SAEs in IFV HAs is short. With a larger number of recorded SAEs in viral surface proteins, machine learning could be implemented to reveal other useful parameters for sequon prediction, or assign weights to existing parameters to enhance prediction accuracy. One potentially interesting parameter not explored in this study is the frequency of near-sequons in IFVs circulating worldwide at a given time. Near-sequon frequencies were reported in Table 4, though it is unclear how these may impact the likelihood of a given sequon emerging. Once further refined, future sequon prediction programs like FSF could be used in synergy with existing algorithms to enhance the accuracy of antigenic escape modeling and help guide the generation of effective vaccines and monoclonal antibody therapies.

## Acknowledgments

The authors would like to thank Dr. Presgraves and Dr. Anderson of the University of Rochester for their helpful suggestions.

## References

1. Newby ML, Allen JD, Crispin M. Influence of glycosylation on the immunogenicity and antigenicity of viral immunogens. Biotechnol Adv. 2024;70:108283. doi:10.1016/j.biotechadv.2023.108283.

2. Zhang XL, Qu H. The Role of Glycosylation in Infectious Diseases. Adv Exp Med Biol. 2021;1325:219–237. doi:10.1007/978-3-030-70115-411.

3. Reis CA, Tauber R, Blanchard V. Glycosylation is a key in SARS-CoV-2 infection. J Mol Med (Berl). 2021;99(8):1023–1031. doi:10.1007/s00109-021-02092-0.

4. Chawla H, Fadda E, Crispin M. Principles of SARS-CoV-2 glycosylation. Curr Opin Struct Biol. 2022;75:102402. doi:10.1016/j.sbi.2022.102402.

5. Idris F, Muharram SH, Diah S. Glycosylation of dengue virus glycoproteins and their interactions with carbohydrate receptors: possible targets for antiviral therapy. Arch Virol. 2016;161(7):1751–60. doi:10.1007/s00705-016-2855-2.

6. Gorzkiewicz M, Cramer J, Xu HC, Lang PA. The role of glycosylation patterns of viral glycoproteins and cell entry receptors in arenavirus infection. Biomed Pharmacother. 2023;166:115196. doi:10.1016/j.biopha.2023.115196.

7. Doud MB, Lee JM, Bloom JD. How single mutations affect viral escape from broad and narrow antibodies to H1 influenza hemagglutinin. Nat Commun. 2018;9(1):1386. doi:10.1038/s41467-018-03665-3.

8. Schon K, Lepenies B, Goyette-Desjardins G. Impact of Protein Glycosylation on the Design of Viral Vaccines. Adv Biochem Eng Biotechnol. 2021;175:319–354. doi:10.1007/102020132.

9. Ozdilek A, Avci FY. Glycosylation as a key parameter in the design of nucleic acid vaccines. Curr Opin Struct Biol. 2022;73:102348. doi:10.1016/j.sbi.2022.102348.

10. Hariharan V, Kane RS. Glycosylation as a tool for rational vaccine design. Biotechnol Bioeng. 2020;117(8):2556–2570. doi:10.1002/bit.27361.

11. Nilchan N, Kraivong R, Luangaram P, Phungsom A, Tantiwatcharakunthon M, Traewachiwiphak S, et al. An Engineered N-Glycosylated Dengue Envelope Protein Domain III Facilitates Epitope-Directed Selection of Potently Neutralizing and Minimally Enhancing Antibodies. ACS Infect Dis. 2024;doi:10.1021/acsinfecdis.4c00058.

12. Martina CE, Crowe J J E, Meiler J. Glycan masking in vaccine design: Targets, immunogens and applications. Front Immunol. 2023;14:1126034. doi:10.3389/fimmu.2023.1126034.

13. Carnell GW, Billmeier M, Vishwanath S, Suau Sans M, Wein H, George CL, et al. Glycan masking of a non-neutralising epitope enhances neutralising antibodies targeting the RBD of SARS-CoV-2 and its variants. Front Immunol. 2023;14:1118523. doi:10.3389/fimmu.2023.1118523.

14. Chuang GY, Boyington JC, Joyce MG, Zhu J, Nabel GJ, Kwong PD, et al. Computational prediction of N-linked glycosylation incorporating structural properties and patterns. Bioinformatics. 2012;28(17):2249–55. doi:10.1093/bioinformatics/bts426.

15. Kornfeld R, Kornfeld S. Assembly of asparagine-linked oligosaccharides. Annu Rev Biochem. 1985;54:631–64. doi:10.1146/annurev.bi.54.070185.003215.

16. Feng T, Zhang J, Chen Z, Pan W, Chen Z, Yan Y, et al. Glycosylation of viral proteins: Implication in virus-host interaction and virulence. Virulence. 2022;13(1):670–683. doi:10.1080/21505594.2022.2060464.

17. Lavie M, Hanoulle X, Dubuisson J. Glycan Shielding and Modulation of Hepatitis C Virus Neutralizing Antibodies. Front Immunol. 2018;9:910. doi:10.3389/fimmu.2018.00910.

18. Grant OC, Montgomery D, Ito K, Woods RJ. Analysis of the SARS-CoV-2 spike protein glycan shield reveals implications for immune recognition. Sci Rep. 2020;10(1):14991. doi:10.1038/s41598-020-71748-7.

19. Gomez Lorenzo MM, Fenton MJ. Immunobiology of influenza vaccines. Chest. 2013;143(2):502–510. doi:10.1378/chest.12-1711.

20. Rodpothong P, Auewarakul P. Viral evolution and transmission effectiveness. World J Virol. 2012;1(5):131–4. doi:10.5501/wjv.v1.i5.131.

21. Tate MD, Job ER, Deng YM, Gunalan V, Maurer-Stroh S, Reading PC. Playing hide and seek: how glycosylation of the influenza virus hemagglutinin can modulate the immune response to infection. Viruses. 2014;6(3):1294–316. doi:10.3390/v6031294.

22. Chen W, Zhong Y, Qin Y, Sun S, Li Z. The evolutionary pattern of glycosylation sites in influenza virus (H5N1) hemagglutinin and neuraminidase. PloS one. 2012;7(11):e49224.

23. Sun S, Wang Q, Zhao F, Chen W, Li Z. Glycosylation site alteration in the evolution of influenza A (H1N1) viruses. PloS one. 2011;6(7):e22844.

24. An Y, Parsons LM, Jankowska E, Melnyk D, Joshi M, Cipollo JF. ¡i¿N¡/i¿-Glycosylation of Seasonal Influenza Vaccine Hemagglutinins: Implication for Potency Testing and Immune Processing. Journal of Virology. 2019;93(2):10.1128/jvi.01693–18. doi:doi:10.1128/jvi.01693-18.

25. Altman MO, Angel M, Kosik I, Trovao NS, Zost SJ, Gibbs JS, et al. Human Influenza A Virus Hemagglutinin Glycan Evolution Follows a Temporal Pattern to a Glycan Limit. mBio. 2019;10(2). doi:10.1128/mBio.00204-19.

26. Oracle Corporation. type [; 2023]Available from: https://www.oracle.com/java/technologies/javase/jdk21-archive-downloads.html.

27. Fraczkiewicz R, Braun W. Exact and Efficient Analytical Calculation of the Accessible Surface Areas and Their Gradients for Macromolecules. Sealy Center for Structural Biology, University of Texas Medical Branch. 1997;.

28. Tamura K, Stecher G, Kumar S. MEGA11: Molecular Evolutionary Genetics Analysis Version 11. Mol Biol Evol. 2021;38(7):3022–3027. doi:10.1093/molbev/msab120.

29. Le SQ, Gascuel O. An improved general amino acid replacement matrix. Mol Biol Evol. 2008;25(7):1307–20. doi:10.1093/molbev/msn067.

30. Jumper J, Evans R, Pritzel A, Green T, Figurnov M, Ronneberger O, et al. Highly accurate protein structure prediction with AlphaFold. Nature. 2021;596(7873):583–589. doi:10.1038/s41586-021-03819-2.

31. Mirdita M, Schutze K, Moriwaki Y, Heo L, Ovchinnikov S, Steinegger M. ColabFold: making protein folding accessible to all. Nat Methods. 2022;19(6):679–682. doi:10.1038/s41592-022-01488-1.

32. Fraczkiewicz R, Braun W. Exact and efficient analytical calculation of the accessible surface areas and their gradients for macromolecules. Journal of Computational Chemistry. 1998;19(3):319–333. doi:Doi 10.1002/(Sici)1096-987x(199802)19:3¡319::Aid-Jcc6¿3.3.Co;2-3.

33. GETAREA;.

34. Edgar RC. MUSCLE: multiple sequence alignment with high accuracy and high throughput. Nucleic Acids Res. 2004;32(5):1792–7. doi:10.1093/nar/gkh340.

35. Olson RD, Assaf R, Brettin T, Conrad N, Cucinell C, Davis JJ, et al. Introducing the Bacterial and Viral Bioinformatics Resource Center (BV-BRC): a resource combining PATRIC, IRD and ViPR. Nucleic Acids Res. 2023;51(D1):D678–D689. doi:10.1093/nar/gkac1003.

36. Shu Y, McCauley J. GISAID: Global initiative on sharing all influenza data - from vision to reality. Euro Surveill. 2017;22(13). doi:10.2807/1560-7917.ES.2017.22.13.30494.

37. Zhang Y, Aevermann BD, Anderson TK, Burke DF, Dauphin G, Gu Z, et al. Influenza Research Database: An integrated bioinformatics resource for influenza virus research. Nucleic Acids Res. 2017;45(D1):D466–D474. doi:10.1093/nar/gkw857.

38. Bogner P, Capua I, Lipman DJ, Cox NJ, et al. A global initiative on sharing avian flu data. Nature. 2006;442(7106):981–981. doi:10.1038/442981a.

39. Boni MF. Vaccination and antigenic drift in influenza. Vaccine. 2008;26 Suppl 3(Suppl 3):C8–14. doi:10.1016/j.vaccine.2008.04.011.

40. Jones S, Nelson-Sathi S, Wang Y, Prasad R, Rayen S, Nandel V, et al. Evolutionary, genetic, structural characterization and its functional implications for the influenza A (H1N1) infection outbreak in India from 2009 to 2017. Sci Rep. 2019;9(1):14690. doi:10.1038/s41598-019-51097-w.

41. Chen Z, Bancej C, Lee L, Champredon D. Publisher Correction: Antigenic drift and epidemiological severity of seasonal influenza in Canada. Sci Rep. 2024;14(1):2952. doi:10.1038/s41598-024-53476-4.

42. Schafer JR, Kawaoka Y, Bean WJ, Suss J, Senne D, Webster RG. Origin of the pandemic 1957 H2 influenza A virus and the persistence of its possible progenitors in the avian reservoir. Virology. 1993;194(2):781–8. doi:10.1006/viro.1993.1319.

43. Wertheim JO. The re-emergence of H1N1 influenza virus in 1977: a cautionary tale for estimating divergence times using biologically unrealistic sampling dates. PLoS One. 2010;5(6):e11184. doi:10.1371/journal.pone.0011184.

44. Rozo M, Gronvall GK. The Reemergent 1977 H1N1 Strain and the Gain-of-Function Debate. mBio. 2015;6(4). doi:10.1128/mBio.01013-15.

45. Otake S, Yoshida M, Dee S. A Review of Swine Breeding Herd Biosecurity in the United States to Prevent Virus Entry Using Porcine Reproductive and Respiratory Syndrome Virus as a Model Pathogen. Animals (Basel). 2024;14(18). doi:10.3390/ani14182694.

46. Jones J, Krag SS, Betenbaugh MJ. Controlling N-linked glycan site occupancy. Biochim Biophys Acta. 2005;1726(2):121–37. doi:10.1016/j.bbagen.2005.07.003.

47. Gupta R, Brunak S. Prediction of glycosylation across the human proteome and the correlation to protein function. Pac Symp Biocomput. 2002; p. 310–22.

48. Pitti T, Chen CT, Lin HN, Choong WK, Hsu WL, Sung TY. N-GlyDE: a two-stage N-linked glycosylation site prediction incorporating gapped dipeptides and pattern-based encoding. Sci Rep. 2019;9(1):15975. doi:10.1038/s41598-019-52341-z.

49. Hou X, Wang Y, Bu D, Wang Y, Sun S. EMNGly: predicting N-linked glycosylation sites using the language models for feature extraction. Bioinformatics. 2023;39(11). doi:10.1093/bioinformatics/btad650.

50. Pakhrin SC, Pokharel S, Aoki-Kinoshita KF, Beck MR, Dam TK, Caragea D, et al. LMNglyPred: prediction of human N-linked glycosylation sites using embeddings from a pre-trained protein language model. Glycobiology. 2023;33(5):411–422. doi:10.1093/glycob/cwad033.

51. Pakhrin SC, Aoki-Kinoshita KF, Caragea D, Kc DB. DeepNGlyPred: A Deep Neural Network-Based Approach for Human N-Linked Glycosylation Site Prediction. Molecules. 2021;26(23). doi:10.3390/molecules26237314.

52. Wagh K, Hahn BH, Korber B. Hitting the sweet spot: exploiting HIV-1 glycan shield for induction of broadly neutralizing antibodies. Curr Opin HIV AIDS. 2020;15(5):267–274. doi:10.1097/COH.0000000000000639.

