## Supplementary material for "Future Sequon Finder - A novel approach for predicting future N-linked glycosylation sequons on viral surface proteins": AminoAcid.java

```
class AminoAcid {
    private String aa;
    private String nucleotides;
    private String nucleotideOrder;
    private int aaOrder;

    public AminoAcid(String aa, String nucleotides, String nucleotideOrder, int aaOrder) {
        this.aa = aa;
        this.nucleotides = nucleotides;
        this.nucleotideOrder = nucleotideOrder;
        this.aaOrder = aaOrder;
    }

    public String getAminoAcid() {
        return aa;
    }

    public String getNucleotides() {
        return nucleotides;
    }

    public String getNucleotideOrder() {
        return nucleotideOrder;
    }

    public int getAminoAcidOrder() {
        return aaOrder;
    }


}
```
