## Supplementary material for "Future Sequon Finder - A novel approach for predicting future N-linked glycosylation sequons on viral surface proteins": GetAreaAminoAcid.java

```
public class GetAreaAminoAcid {
    private String singleLetterIdentifier;
    private int positionIdentifier;
    private double exposureRatio;
    private boolean isOut;

    public GetAreaAminoAcid(String singleLetterIdentifier, int positionIdentifier, double exposureRatio, boolean isOut) {
        this.singleLetterIdentifier = singleLetterIdentifier;
        this.positionIdentifier = positionIdentifier;
        this.exposureRatio = exposureRatio;
        this.isOut = isOut;
    }

    public String getSingleLetterIdentifier() {
        return singleLetterIdentifier;
    }

    public int getPositionIdentifier() {
        return positionIdentifier;
    }

    public double getExposureRatio() {
        return exposureRatio;
    }

    public boolean isOut() {
        return isOut;
    }
}
```
