## Supplementary material for "Future Sequon Finder - A novel approach for predicting future N-linked glycosylation sequons on viral surface proteins": GetAreaInputProcessor.java

```
import java.util.ArrayList;
import java.util.List;

public class GetAreaInputProcessor {

    public static List<GetAreaAminoAcid> processInput(String input) {
        List<GetAreaAminoAcid> areaAminoAcids = new ArrayList<>();

        String[] lines = input.split("\n");
        for (String line : lines) {

            line = line.trim();

            String[] tokens = line.split("\\s+");

            if (tokens.length >= 7) {
                String aaName = tokens[0];
                int position = Integer.parseInt(tokens[1]);
                double exposureRatio = Double.parseDouble(tokens[6]);
                boolean isOut = tokens.length > 7 && tokens[7].equals("o");

                String singleLetterIdentifier = convertToSingleLetter(aaName);

                GetAreaAminoAcid aminoAcid = new GetAreaAminoAcid(singleLetterIdentifier, position, exposureRatio, isOut);
                areaAminoAcids.add(aminoAcid);
            }
        }

        return areaAminoAcids;
    }

    private static String convertToSingleLetter(String threeLetterIdentifier) {
        switch (threeLetterIdentifier) {
            case "ALA":
                return "A";
            case "ARG":
                return "R";
            case "ASN":
                return "N";
            case "ASP":
                return "D";
            case "CYS":
                return "C";
            case "GLN":
                return "Q";
            case "GLU":
                return "E";
            case "GLY":
                return "G";
            case "HIS":
                return "H";
            case "ILE":
                return "I";
            case "LEU":
                return "L";
            case "LYS":
                return "K";
            case "MET":
                return "M";
            case "PHE":
                return "F";
            case "PRO":
                return "P";
            case "SER":
                return "S";
            case "THR":
                return "T";
            case "TRP":
                return "W";
            case "TYR":
                return "Y";
            case "VAL":
                return "V";
            default:
                return "?";
        }
    }
}
```
