## Supplementary material for "Future Sequon Finder - A novel approach for predicting future N-linked glycosylation sequons on viral surface proteins": Main.java

```
//TIP To <b>Run</b> code, press <shortcut actionId="Run"/> or
// click the <icon src="AllIcons.Actions.Execute"/> icon in the gutter.
import javax.swing.*;
import java.awt.*;
import java.awt.event.ActionEvent;
import java.awt.event.ActionListener;
import java.util.ArrayList;
import java.util.List;
import javax.swing.border.EmptyBorder;
import java.awt.event.ItemEvent;
import java.awt.event.ItemListener;
import java.util.Map;
import java.util.HashMap;
import java.util.stream.Collectors;

public class Main {
    private static JTextArea outputTextArea;
    private static JTextArea inputField;
    private static JTextArea areaInputField;

    private static JLabel inputLabel;
    private static JTextArea megaInputField;
    private static List<String> veryNearSequons;
    private static List<String> ultraNearSequons;
    private static List<GetAreaAminoAcid> getAreaAminoAcids;
    private static List<String> sequonsWithExposedSurface;
    private static List<String> nearSequonsWithExposedSurface;
    private static List<String> veryNearSequonsWithExposedSurface;
    private static List<String> ultraNearSequonsWithExposedSurface;
    private static JCheckBox checkBox1;
    private static JCheckBox checkBox2;
    private static JCheckBox checkBox3;
    private static JCheckBox checkBox4;
    private static JCheckBox checkBox5;
    private static JCheckBox checkBox6;
    private static JCheckBox checkBox7;
    private static JCheckBox checkBox8;
    private static JCheckBox checkBox9;
    private static JCheckBox checkBox10;
    private static JCheckBox checkBox11;
    private static JCheckBox checkBox12;

    private static JCheckBox singleSequenceAnalysisCheckBox;
    private static JCheckBox multipleSequenceAnalysisCheckBox;

    public static void main(String[] args) {
        JFrame frame = new JFrame("Future Sequon Finder v1.1");
        frame.setDefaultCloseOperation(JFrame.EXIT_ON_CLOSE);
        frame.setSize(1850, 1000);

        int marginSize = 15;
        ((JComponent) frame.getContentPane()).setBorder(new EmptyBorder(marginSize, marginSize, marginSize, marginSize));
        frame.setLayout(new BorderLayout());


        // Add the analysis mode panel to the existing title label panel or create a new panel for both
        JPanel titleAndAnalysisModePanel = new JPanel(new BorderLayout());
        JLabel titleLabel = new JLabel("Future Sequon Finder - Zand Lab", SwingConstants.CENTER);
        titleLabel.setFont(new Font("Arial", Font.BOLD, 20));
        titleAndAnalysisModePanel.add(titleLabel, BorderLayout.CENTER);

        frame.add(titleAndAnalysisModePanel, BorderLayout.NORTH);

        outputTextArea = new JTextArea();
        outputTextArea.setFont(new Font("Monospaced", Font.PLAIN, 12));
        outputTextArea.setEditable(false);
        JScrollPane scrollPane = new JScrollPane(outputTextArea);
        frame.add(scrollPane, BorderLayout.CENTER);

        JPanel inputPanel = new JPanel();
        inputPanel.setLayout(new BoxLayout(inputPanel, BoxLayout.Y_AXIS));

        JPanel checkboxesPanel = new JPanel(new GridLayout(4,3));

        checkBox1 = new JCheckBox("Display existing sequons");
        checkBox2 = new JCheckBox("Display near-sequons");
        checkBox3 = new JCheckBox("Display very near-sequons");
        checkBox4 = new JCheckBox("Display ultra near-sequons");
        checkBox5 = new JCheckBox("Use GetArea Surface Accessibility feature");
        checkBox6 = new JCheckBox("Display near-sequons with exposed surfaces");
        checkBox7 = new JCheckBox("Display very near-sequons with exposed surfaces");
        checkBox8 = new JCheckBox("Display ultra near-sequons with exposed surfaces");
        checkBox9 = new JCheckBox("Use MEGA mutation feature");
        checkBox10 = new JCheckBox("Sort all near-sequons");
        checkBox11 = new JCheckBox("Sort all very near-sequons");
        checkBox12 = new JCheckBox("Sort all ultra near-sequons");

        checkboxesPanel.add(checkBox1);
        checkboxesPanel.add(checkBox5);
        checkboxesPanel.add(checkBox9);

        checkboxesPanel.add(checkBox2);
        checkboxesPanel.add(checkBox6);
        checkboxesPanel.add(checkBox10);

        checkboxesPanel.add(checkBox3);
        checkboxesPanel.add(checkBox7);
        checkboxesPanel.add(checkBox11);

        checkboxesPanel.add(checkBox4);
        checkboxesPanel.add(checkBox8);
        checkboxesPanel.add(checkBox12);

        inputPanel.add(checkboxesPanel, BorderLayout.NORTH);

        JPanel firstInputPanel = new JPanel(new BorderLayout());
        inputLabel = new JLabel("Input nucleotide sequence here:   ", SwingConstants.LEFT);
        inputField = new JTextArea(10, 100);
        inputField.setLineWrap(true);
        inputField.setWrapStyleWord(true);
        JScrollPane firstScrollPane = new JScrollPane(inputField);
        firstInputPanel.add(inputLabel, BorderLayout.WEST);
        firstInputPanel.add(firstScrollPane, BorderLayout.CENTER);
        inputPanel.add(firstInputPanel);

        JPanel secondInputPanel = new JPanel(new BorderLayout());
        JLabel areaInputLabel = new JLabel("Input 'GetArea' surface accessibility result:   ", SwingConstants.LEFT);
        areaInputField = new JTextArea(10, 100);
        areaInputField.setLineWrap(true);
        areaInputField.setWrapStyleWord(true);
        JScrollPane secondScrollPane = new JScrollPane(areaInputField);
        secondInputPanel.add(areaInputLabel, BorderLayout.WEST);
        secondInputPanel.add(secondScrollPane, BorderLayout.CENTER);
        inputPanel.add(secondInputPanel);

        JPanel thirdInputPanel = new JPanel(new BorderLayout());
        JLabel megaInputLabel = new JLabel("Input 'MEGA' mutation frequency result:   ", SwingConstants.LEFT);
        megaInputField = new JTextArea(10, 100);
        megaInputField.setLineWrap(true);
        megaInputField.setWrapStyleWord(true);
        JScrollPane thirdScrollPane = new JScrollPane(megaInputField);
        thirdInputPanel.add(megaInputLabel, BorderLayout.WEST);
        thirdInputPanel.add(thirdScrollPane, BorderLayout.CENTER);
        inputPanel.add(thirdInputPanel);


        JPanel buttonsPanel = new JPanel();
        buttonsPanel.setLayout(new FlowLayout(FlowLayout.RIGHT));
        JButton submitButton = new JButton("Submit");
        submitButton.addActionListener(new ActionListener() {
            @Override
            public void actionPerformed(ActionEvent e) {
                processInput();
            }
        });
        buttonsPanel.add(submitButton);

        JButton resetButton = new JButton("Reset");
        resetButton.addActionListener(new ActionListener() {
            @Override
            public void actionPerformed(ActionEvent e) {
                resetProgram();
            }
        });
        buttonsPanel.add(resetButton);

        JButton helpButton = new JButton("Help");
        helpButton.addActionListener(new ActionListener() {
            @Override
            public void actionPerformed(ActionEvent e) {

                outputTextArea.setText(" Nucleotide Based Glycan Finder is a program that processes a nucleotide sequence from the user and converts it into an amino acid sequence."
                        + "\n\n The resulting amino acid sequence is then scanned for N-linked glycosylation sequons, and sites that may become glycosylation sequons using the known N-linked glycosylation sequon definition (N - !P - S/T)."
                        + "\n\n These sites will be sorted into the following categories:"
                        + "\n\n\n    Near-sequon       - A site that may become an N-linked glycosylation sequon if a single amino acid is changed"
                        + "\n\n    Very near-sequon  - A site that may become an N-linked glycosylation sequon if a single nucleotide is changed"
                        + "\n\n    Ultra near-sequon - A site that may become an N-linked glycosylation sequon if a single nucleotide is changed, and the resultant change does not alter the charge or polarity of the amino acid"
                        + "\n\n\n The 'GetArea' surface accessability feature allows users to pair their nucleic acid sequence with surface accessability results from https://curie.utmb.edu/getarea.html to better select for near glycosylation sequons."
                        + "\n\n When using this feature, it is critical that the amino acid sequence generated from the nucleotide input matches the amino acid sequence AND position numbers of the PDB file provided to the GetArea program."
                        + "\n\n When using GetArea, use the default probe radius of 1.4 angstrom."
                        + "\n\n To input the results from GetArea into the Nucleotide Based Glycan Finder, simply copy and paste the query results from GetArea, starting with the first amino acid, into the associated input box in NBGF after pasting the corresponding nucleotide sequence in the box above."
                        + "\n\n    Note: Do not include the 'Probe radius' or any of the columb titles in the input for your GetArea results, as this will result in an error."
                        + "\n\n\n If all amino acids in the GetArea result allign with the amino acids generated from the nucleotide sequence, the sidechain surface exposure ratio will be saved to each exposed amino acid."
                        + "\n\n Any amino acid associated with the asparagine (N) site of a near-sequon that has a side chain exposure ratio exceeding 15% will be marked as exposed."
                        + "\n\n\n Use the check boxes to select the lists that are to be printed once the 'Submit' button is pressed."
                        + "\n\n Press the 'Reset' button at any time to reset the program to its launch state.");
            }
        });
        buttonsPanel.add(helpButton);


        JPanel inputAndCheckboxesPanel = new JPanel(new BorderLayout());
        inputAndCheckboxesPanel.add(inputPanel, BorderLayout.CENTER);
        inputAndCheckboxesPanel.add(buttonsPanel, BorderLayout.SOUTH);


        checkBox6.setEnabled(false);
        checkBox7.setEnabled(false);
        checkBox8.setEnabled(false);
        areaInputLabel.setEnabled(false);
        areaInputField.setEnabled(false);

        checkBox5.addItemListener(new ItemListener() {
            @Override
            public void itemStateChanged(ItemEvent e) {

                boolean useGetAreaFeature = checkBox5.isSelected();

                checkBox6.setEnabled(useGetAreaFeature);
                checkBox7.setEnabled(useGetAreaFeature);
                checkBox8.setEnabled(useGetAreaFeature);

                areaInputField.setEnabled(useGetAreaFeature);
                areaInputLabel.setEnabled(useGetAreaFeature);

                if (!useGetAreaFeature) {
                    checkBox6.setSelected(false);
                    checkBox7.setSelected(false);
                    checkBox8.setSelected(false);
                    areaInputField.setText("");
                }
            }
        });

        checkBox10.setEnabled(false);
        checkBox11.setEnabled(false);
        checkBox12.setEnabled(false);
        megaInputLabel.setEnabled(false);
        megaInputField.setEnabled(false);

        checkBox9.addItemListener(new ItemListener() {
            @Override
            public void itemStateChanged(ItemEvent e) {

                boolean useGetAreaFeature = checkBox9.isSelected();

                checkBox10.setEnabled(useGetAreaFeature);
                checkBox11.setEnabled(useGetAreaFeature);
                checkBox12.setEnabled(useGetAreaFeature);

                megaInputField.setEnabled(useGetAreaFeature);
                megaInputLabel.setEnabled(useGetAreaFeature);

                if (!useGetAreaFeature) {
                    checkBox10.setSelected(false);
                    checkBox11.setSelected(false);
                    checkBox12.setSelected(false);
                    megaInputField.setText("");
                }
            }
        });

        frame.add(inputAndCheckboxesPanel, BorderLayout.SOUTH);


        veryNearSequons = new ArrayList<>();
        ultraNearSequons = new ArrayList<>();
        sequonsWithExposedSurface = new ArrayList<>();
        nearSequonsWithExposedSurface = new ArrayList<>();
        veryNearSequonsWithExposedSurface = new ArrayList<>();
        ultraNearSequonsWithExposedSurface = new ArrayList<>();
        getAreaAminoAcids = new ArrayList<>();


        frame.setVisible(true);

        outputTextArea.append("Welcome to the Nucleotide Based Glycan Finder! Please input a nucleotide sequence below. The program will then convert it into an amino acid sequence and check for N-linked glycosylation sequons as well as sites that may easily be transformed into sequons."
                + "\n\n\nFor more information about the program, press the 'Help' button.");
    }


    private static void processInput() {
        String inputSequence;
        String inputSequenceUnrefined = inputField.getText();


        inputSequence = inputSequenceUnrefined.replaceAll("\n", "")
                .replaceAll("\\d", "")
                .replaceAll(" ", "")
                .toUpperCase();

        int invalidPosition = findInvalidPosition(inputSequence);

        if (invalidPosition != -1) {
            char invalidChar = inputSequence.charAt(invalidPosition);
            System.out.println("Invalid character '" + invalidChar + "' found in nucleotide sequence at position " + (invalidPosition + 1));

            outputTextArea.setText("");
            outputTextArea.append("Error: Unacceptable character '" + invalidChar + "' entered in nucleotide sequence input at position " + (invalidPosition + 1) + ". Input sequence should consist of only 'A', 'T', 'G', and 'C'. Spaces, indentations, numbers, and lowercase nucleotides are acceptable as well and will be deleted upon entry.\n");
            return;
        }


        String getAreaInput = areaInputField.getText();
        getAreaAminoAcids = GetAreaInputProcessor.processInput(getAreaInput);

        List<AminoAcid> aminoAcids = NucleotideToAminoAcid.convertToAminoAcids(inputSequence);
        StringBuilder aminoAcidSequence = new StringBuilder("Amino acid sequence: ");


        for (AminoAcid aa : aminoAcids) {
            aminoAcidSequence.append(aa.getAminoAcid());
        }


        List<String> sequons = SequonScanner.findSequons(aminoAcids);

        List<String> nearSequons = NearSequonScanner.findNearSequons(aminoAcids, sequons);

        List<String> nearSequonsWithExposedSurface = new ArrayList<>();

        List<String> veryNearSequonsWithExposedSurface = new ArrayList<>();

        List<String> ultraNearSequonsWithExposedSurface = new ArrayList<>();


        if (checkBox5.isSelected()) {
            Map<Integer, Integer> adjustedPositionMap = createAdjustedPositionMap(aminoAcids);
            for (int i = 0; i < sequons.size(); i++) {
                String sequon = sequons.get(i);
                int aminoAcidPosition = extractAminoAcidPosition(sequon);
                int adjustedPosition = adjustedPositionMap.getOrDefault(aminoAcidPosition, -1);

                GetAreaAminoAcid matchingGetAreaAminoAcid = getAreaAminoAcids.stream()
                        .filter(aa -> aa.getPositionIdentifier() == adjustedPosition)
                        .findFirst()
                        .orElse(null);

                if (matchingGetAreaAminoAcid != null) {
                    AminoAcid originalAA = aminoAcids.stream()
                            .filter(aa -> aa.getAminoAcidOrder() == aminoAcidPosition)
                            .findFirst()
                            .orElse(null);
                    if (originalAA != null && !matchingGetAreaAminoAcid.getSingleLetterIdentifier().equals(originalAA.getAminoAcid())) {
                        outputTextArea.setText("");
                        outputTextArea.append("Error: Mismatch between GetArea amino acid sequence and amino acid sequence generated from user input detected - Position: " + aminoAcidPosition + " AA: " + matchingGetAreaAminoAcid.getSingleLetterIdentifier() + ", press 'Help' for more assistance.");
                        return;
                    }
                    double exposureRatio = matchingGetAreaAminoAcid.getExposureRatio();
                    String updatedSequon = sequon + ", Side Chain Exposure Ratio: " + (exposureRatio > 0 ? exposureRatio + "%" : "NA");
                    sequons.set(i, updatedSequon);
                    if (exposureRatio > 15) {
                        sequonsWithExposedSurface.add(updatedSequon);
                    }
                } else {
                    sequons.set(i, sequon + ", Side Chain Exposure Ratio: NA");
                }
            }
        }

        if (checkBox5.isSelected()) {
            Map<Integer, Integer> adjustedPositionMap = createAdjustedPositionMap(aminoAcids);
            for (int i = 0; i < nearSequons.size(); i++) {
                String nearSequon = nearSequons.get(i);
                int aminoAcidPosition = extractAminoAcidPosition(nearSequon);
                int adjustedPosition = adjustedPositionMap.getOrDefault(aminoAcidPosition, -1);

                GetAreaAminoAcid matchingGetAreaAminoAcid = getAreaAminoAcids.stream()
                        .filter(aa -> aa.getPositionIdentifier() == adjustedPosition)
                        .findFirst()
                        .orElse(null);

                if (matchingGetAreaAminoAcid != null) {
                    AminoAcid originalAA = aminoAcids.get(aminoAcidPosition - 1);
                    if (!matchingGetAreaAminoAcid.getSingleLetterIdentifier().equals(originalAA.getAminoAcid())) {
                        outputTextArea.setText("");
                        outputTextArea.append("Error: Mismatch between GetArea amino acid sequence and amino acid sequence generated from user input detected - Position: " + aminoAcidPosition + " AA: " + matchingGetAreaAminoAcid.getSingleLetterIdentifier() + ", press 'Help' for more assistance.");
                        return;
                    }
                    double exposureRatio = matchingGetAreaAminoAcid.getExposureRatio();
                    String updatedNearSequon = nearSequon + ", Side Chain Exposure Ratio: " + (exposureRatio > 0 ? exposureRatio + "%" : "NA");
                    nearSequons.set(i, updatedNearSequon);
                    if (exposureRatio > 15) {
                        nearSequonsWithExposedSurface.add(updatedNearSequon);
                    }
                } else {
                    nearSequons.set(i, nearSequon + ", Side Chain Exposure Ratio: NA");
                }
            }
        }

        for (String nearSequon : nearSequonsWithExposedSurface) {
            boolean singleNucleotideChange = nearSequon.contains("Single nucleotide change: true");
            if (singleNucleotideChange) {
                veryNearSequonsWithExposedSurface.add(nearSequon);
            }

            boolean ultraNearSequonCondition = nearSequon.contains("Single nucleotide change: true") && nearSequon.contains("Same charge & polarity: true");
            if (ultraNearSequonCondition) {
                ultraNearSequonsWithExposedSurface.add(nearSequon);
            }
        }


        for (String nearSequon : nearSequons) {
            boolean singleNucleotideChange = nearSequon.contains("Single nucleotide change: true");
            if (singleNucleotideChange) {
                veryNearSequons.add(nearSequon);
            }
        }

        for (String nearSequon : nearSequons) {
            boolean singleNucleotideChange = nearSequon.contains("Single nucleotide change: true") && nearSequon.contains("Same charge & polarity: true");
            if (singleNucleotideChange) {
                ultraNearSequons.add(nearSequon);
            }


        }

        List<String> mutSortedNearSequons = new ArrayList<>();
        List<String> mutSortedVeryNearSequons = new ArrayList<>();
        List<String> mutSortedUltraNearSequons = new ArrayList<>();
        List<String> mutSortedNearSequonsWithExposedSurface = new ArrayList<>();
        List<String> mutSortedVeryNearSequonsWithExposedSurface = new ArrayList<>();
        List<String> mutSortedUltraNearSequonsWithExposedSurface = new ArrayList<>();


        if (checkBox9.isSelected()) {
            String megaInput = megaInputField.getText();
            List<MegaSiteRate> megaSiteRates = MegaInputProcessor.processInput(megaInput);
            mutSortedNearSequons = processAndSortNearSequons(nearSequons, megaSiteRates);
            mutSortedVeryNearSequons = processAndSortNearSequons(veryNearSequons, megaSiteRates);
            mutSortedUltraNearSequons = processAndSortNearSequons(ultraNearSequons, megaSiteRates);
            mutSortedNearSequonsWithExposedSurface = processAndSortNearSequons(nearSequonsWithExposedSurface, megaSiteRates);
            mutSortedVeryNearSequonsWithExposedSurface = processAndSortNearSequons(veryNearSequonsWithExposedSurface, megaSiteRates);
            mutSortedUltraNearSequonsWithExposedSurface = processAndSortNearSequons(ultraNearSequonsWithExposedSurface, megaSiteRates);
        }


        formatAndDisplayOutput(
                aminoAcids,
                sequons,
                nearSequons,
                veryNearSequons,
                ultraNearSequons,
                nearSequonsWithExposedSurface,
                veryNearSequonsWithExposedSurface,
                ultraNearSequonsWithExposedSurface,
                mutSortedNearSequons,
                mutSortedVeryNearSequons,
                mutSortedUltraNearSequons,
                mutSortedNearSequonsWithExposedSurface,
                mutSortedVeryNearSequonsWithExposedSurface,
                mutSortedUltraNearSequonsWithExposedSurface);
    }

    private static int findInvalidPosition(String inputSequence) {
        for (int i = 0; i < inputSequence.length(); i++) {
            char nucleotide = inputSequence.charAt(i);

            if (nucleotide != 'A' && nucleotide != 'T' && nucleotide != 'G' && nucleotide != 'C' && nucleotide != ' ' && nucleotide != '-') {
                return i;
            }
        }
        return -1;
    }

    private static void formatAndDisplayOutput(
            List<AminoAcid> aminoAcids,
            List<String> sequons,
            List<String> nearSequons,
            List<String> veryNearSequons,
            List<String> ultraNearSequons,
            List<String> nearSequonsWithExposedSurface,
            List<String> veryNearSequonsWithExposedSurface,
            List<String> ultraNearSequonsWithExposedSurface,
            List<String> mutSortedNearSequons,
            List<String> mutSortedVeryNearSequons,
            List<String> mutSortedUltraNearSequons,
            List<String> mutSortedNearSequonsWithExposedSurface,
            List<String> mutSortedVeryNearSequonsWithExposedSurface,
            List<String> mutSortedUltraNearSequonsWithExposedSurface
    ) {

        String inputSequence;
        String inputSequenceUnrefined = inputField.getText();
        boolean displayExistingSequons = checkBox1.isSelected();
        boolean displayNearSequons = checkBox2.isSelected();
        boolean displayVeryNearSequons = checkBox3.isSelected();
        boolean displayUltraNearSequons = checkBox4.isSelected();

        boolean useSurfaceAccessibility = checkBox5.isSelected();
        boolean displayNearSequonsExposed = checkBox6.isSelected();
        boolean displayVeryNearSequonsExposed = checkBox7.isSelected();
        boolean displayUltraNearSequonsExposed = checkBox8.isSelected();
        boolean useMutSortedSequons = checkBox9.isSelected();
        boolean displayMutSortedNearSequons = checkBox10.isSelected();
        boolean displayMutSortedVeryNearSequons = checkBox11.isSelected();
        boolean displayMutSortedUltraNearSequons = checkBox12.isSelected();

        inputSequence = inputSequenceUnrefined.replaceAll("\n", "")
                .replaceAll("\\d", "")
                .replaceAll(" ", "")
                .toUpperCase();

        outputTextArea.setText("");

        outputTextArea.append("Nucleotide sequence input: " + inputSequence + "\n\n");

        outputTextArea.append("Amino acid sequence: ");
        for (AminoAcid aa : aminoAcids) {
            outputTextArea.append(aa.getAminoAcid());
        }
        outputTextArea.append("\n\n");

        // This ensures displayed sequons and near sequons are in the head region of the H1 HA domain (AA 60-300). Adjust as needed.
        java.util.function.Predicate<String> positionFilter = sequon -> {
            int position = extractAminoAcidPosition(sequon);
            return position > 60 && position < 300;
        };


        if (displayExistingSequons) {
            List<String> filteredSequons = sequons.stream().filter(positionFilter).collect(Collectors.toList());
            outputTextArea.append("List of all existing N-linked glycosylation sequons in amino acid sequence:\n\n");
            displayFormattedList(filteredSequons);
            outputTextArea.append("\nTotal number of sequons found: " + filteredSequons.size() + "\n\n\n\n");
        }

        if (displayNearSequons) {
            List<String> filteredNearSequons = nearSequons.stream().filter(positionFilter).collect(Collectors.toList());
            outputTextArea.append("List of near-sequons, defined as being able to become a sequon if only one amino acid is changed:\n\n");
            displayFormattedList(filteredNearSequons);
            outputTextArea.append("\nTotal number of near-sequons found: " + filteredNearSequons.size() + "\n\n\n\n");
        }

        if (displayVeryNearSequons) {
            List<String> filteredVeryNearSequons = veryNearSequons.stream().filter(positionFilter).collect(Collectors.toList());
            outputTextArea.append("List of very near-sequons, defined as being able to become a sequon if only one nucleotide is changed:\n\n");
            displayFormattedList(filteredVeryNearSequons);
            outputTextArea.append("\nTotal number of very near-sequons found: " + filteredVeryNearSequons.size() + "\n\n\n\n");
        }

        if (displayUltraNearSequons) {
            List<String> filteredUltraNearSequons = ultraNearSequons.stream().filter(positionFilter).collect(Collectors.toList());
            outputTextArea.append("List of ultra near-sequons, defined as a very near-sequon in which the single nucleotide change does not change the charge or polarity of the original AA:\n\n");
            displayFormattedList(filteredUltraNearSequons);
            outputTextArea.append("\nTotal number of ultra near-sequons found: " + filteredUltraNearSequons.size() + "\n\n\n\n\n\n");
        }


        if (useSurfaceAccessibility) {
            if (displayExistingSequons) {
                List<String> filteredSequonsWithExposedSurface = sequonsWithExposedSurface.stream().filter(positionFilter).collect(Collectors.toList());
                outputTextArea.append("List of existing sequons for which the first asparagine residue has a sidechain exposure exceeding 15%:\n\n");
                displayFormattedList(filteredSequonsWithExposedSurface);
                outputTextArea.append("\nTotal number of sequons found with exposed sidechain: " + filteredSequonsWithExposedSurface.size() + "\n\n\n\n");
            }

            if (displayNearSequonsExposed) {
                List<String> filteredNearSequonsWithExposedSurface = nearSequonsWithExposedSurface.stream().filter(positionFilter).collect(Collectors.toList());
                outputTextArea.append("List of near-sequons for which the first AA has a sidechain exposure exceeding 15%: \n\n");
                displayFormattedList(filteredNearSequonsWithExposedSurface);
                outputTextArea.append("\nTotal number of near-Sequons with exposed sidechain: " + filteredNearSequonsWithExposedSurface.size() + "\n\n\n\n");
            }

            if (displayVeryNearSequonsExposed) {
                List<String> filteredVeryNearSequonsWithExposedSurface = veryNearSequonsWithExposedSurface.stream().filter(positionFilter).collect(Collectors.toList());
                outputTextArea.append("List of very near-sequons for which the first AA has a sidechain exposure exceeding 15%:\n\n");
                displayFormattedList(filteredVeryNearSequonsWithExposedSurface);
                outputTextArea.append("\nTotal number of very near-Sequons with exposed sidechain: " + filteredVeryNearSequonsWithExposedSurface.size() + "\n\n\n\n");
            }

            if (displayUltraNearSequonsExposed) {
                List<String> filteredUltraNearSequonsWithExposedSurface = ultraNearSequonsWithExposedSurface.stream().filter(positionFilter).collect(Collectors.toList());
                outputTextArea.append("List of ultra near-sequons for which the first AA has a sidechain exposure exceeding 15%:\n\n");
                displayFormattedList(filteredUltraNearSequonsWithExposedSurface);
                outputTextArea.append("\nTotal number of ultra near-Sequons with exposed sidechain: " + filteredUltraNearSequonsWithExposedSurface.size() + "\n\n\n\n\n\n");
            }
        }

        if (useMutSortedSequons) {
            if (displayMutSortedNearSequons) {
                List<String> filteredMutSortedNearSequons = mutSortedNearSequons.stream().filter(positionFilter).collect(Collectors.toList());
                outputTextArea.append("Sorted near-sequons based on the mutation frequency of the amino acid site that must change (highest to lowest):\n\n");
                displayFormattedList(filteredMutSortedNearSequons);
                outputTextArea.append("\nTotal number of near-Sequons sorted by mutation rate: " + filteredMutSortedNearSequons.size() + "\n\n\n\n");

                if (displayNearSequonsExposed) {
                    List<String> filteredMutSortedNearSequonsWithExposedSurface = mutSortedNearSequonsWithExposedSurface.stream().filter(positionFilter).collect(Collectors.toList());
                    outputTextArea.append("List of near-sequons for which the first AA has a sidechain exposure exceeding 15% sorted by mutation rate: \n\n");
                    displayFormattedList(filteredMutSortedNearSequonsWithExposedSurface);
                    outputTextArea.append("\nTotal number of near-Sequons with exposed sidechain sorted by mutation rate: " + filteredMutSortedNearSequonsWithExposedSurface.size() + "\n\n\n\n");
                }
            }

            if (displayMutSortedVeryNearSequons) {
                List<String> filteredMutSortedVeryNearSequons = mutSortedVeryNearSequons.stream().filter(positionFilter).collect(Collectors.toList());
                outputTextArea.append("Sorted very near-sequons based on the mutation frequency of the amino acid site that must change (highest to lowest):\n\n");
                displayFormattedList(filteredMutSortedVeryNearSequons);
                outputTextArea.append("\nTotal number of very near-Sequons sorted by mutation rate: " + filteredMutSortedVeryNearSequons.size() + "\n\n\n\n");

                if (displayVeryNearSequonsExposed) {
                    List<String> filteredMutSortedVeryNearSequonsWithExposedSurface = mutSortedVeryNearSequonsWithExposedSurface.stream().filter(positionFilter).collect(Collectors.toList());
                    outputTextArea.append("List of very near-sequons for which the first AA has a sidechain exposure exceeding 15% sorted by mutation rate:\n\n");
                    displayFormattedList(filteredMutSortedVeryNearSequonsWithExposedSurface);
                    outputTextArea.append("\nTotal number of very near-Sequons with exposed sidechain sorted by mutation rate: " + filteredMutSortedVeryNearSequonsWithExposedSurface.size() + "\n\n\n\n");
                }
            }

            if (displayMutSortedUltraNearSequons) {
                List<String> filteredMutSortedUltraNearSequons = mutSortedUltraNearSequons.stream().filter(positionFilter).collect(Collectors.toList());
                outputTextArea.append("Sorted ultra near-sequons based on the mutation frequency of the amino acid site that must change (highest to lowest):\n\n");
                displayFormattedList(filteredMutSortedUltraNearSequons);
                outputTextArea.append("\nTotal number of ultra near-Sequons sorted by mutation rate: " + filteredMutSortedUltraNearSequons.size() + "\n\n\n\n");

                if (displayUltraNearSequonsExposed) {
                    List<String> filteredMutSortedUltraNearSequonsWithExposedSurface = mutSortedUltraNearSequonsWithExposedSurface.stream().filter(positionFilter).collect(Collectors.toList());
                    outputTextArea.append("List of ultra near-sequons for which the first AA has a sidechain exposure exceeding 15% sorted by mutation rate:\n\n");
                    displayFormattedList(filteredMutSortedUltraNearSequonsWithExposedSurface);
                    outputTextArea.append("\nTotal number of ultra near-Sequons with exposed sidechain sorted by mutation rate: " + filteredMutSortedUltraNearSequonsWithExposedSurface.size() + "\n\n\n\n");
                }
            }
        }

        veryNearSequons.clear();
        ultraNearSequons.clear();
        sequonsWithExposedSurface.clear();
    }

    private static void displayFormattedList(List<String> list) {
        for (String item : list) {
            String[] parts = item.split(", ");
            for (String part : parts) {
                outputTextArea.append(String.format("%-50s", part));
            }
            outputTextArea.append("\n");
        }
    }

    private static void resetProgram() {

        veryNearSequons.clear();
        ultraNearSequons.clear();
        sequonsWithExposedSurface.clear();

        inputField.setText("");

        areaInputField.setText("");

        megaInputField.setText("");

        outputTextArea.setText("");

        outputTextArea.append("Welcome to the Nucleotide Based Glycan Finder! Please input a nucleotide sequence below. The program will then convert it into an amino acid sequence and check for N-linked glycosylation sequons as well as sites that may easily be transformed into sequons."
                + "\n\n\nFor more information about the program, press the 'Help' button.");
    }


    private static int extractAminoAcidPosition(String nearSequon) {
        java.util.regex.Pattern pattern = java.util.regex.Pattern.compile("at AA position (\\d+):");
        java.util.regex.Matcher matcher = pattern.matcher(nearSequon);

        if (matcher.find()) {
            return Integer.parseInt(matcher.group(1));
        }

        return -1;
    }


    private static Map<Integer, Integer> createAdjustedPositionMap(List<AminoAcid> aminoAcids) {
        Map<Integer, Integer> adjustedPositionMap = new HashMap<>();
        int adjustedPosition = 1;
        for (int i = 0; i < aminoAcids.size(); i++) {
            if (!aminoAcids.get(i).getAminoAcid().equals("-")) {
                adjustedPositionMap.put(i + 1, adjustedPosition++);
            }
        }
        return adjustedPositionMap;
    }

    private static List<String> processAndSortNearSequons(List<String> sequons, List<MegaSiteRate> megaSiteRates) {
        Map<Integer, Double> mutationRateMap = megaSiteRates.stream()
                .collect(Collectors.toMap(MegaSiteRate::getPositionIdentifier, MegaSiteRate::getMutationRate));

        List<String> sortedSequons = new ArrayList<>();
        for (String sequon : sequons) {
            int position = extractAminoAcidPosition(sequon);
            if (position <= 60 || position >= 300) {
                continue;  // Skip sequons outside the AA position range of 60-300 (outside of the H1 head domain)
            }
            String aaIdentifier;
            int offset;
            if (sequon.contains("Mismatched codon (AA1")) {
                aaIdentifier = "AA1";
                offset = 0;
            } else if (sequon.contains("Mismatched codon (AA2")) {
                aaIdentifier = "AA2";
                offset = 1;
            } else {
                aaIdentifier = "AA3";
                offset = 2;
            }
            double mutationRate = mutationRateMap.getOrDefault(position + offset, 0.0);
            String updatedSequon = sequon + ", Mutation rate of site " + (position + offset) + " (" + aaIdentifier + "): " + mutationRate;
            sortedSequons.add(updatedSequon);
        }

        sortedSequons.sort((a, b) -> {
            double rateA = Double.parseDouble(a.substring(a.lastIndexOf(":") + 1));
            double rateB = Double.parseDouble(b.substring(b.lastIndexOf(":") + 1));
            return Double.compare(rateB, rateA);
        });

        // Re-rank the sequons after sorting and filtering
        List<String> rankedSequons = new ArrayList<>();
        int rank = 1;
        for (String sequon : sortedSequons) {
            String rankedSequon = String.format("#%d %s", rank, sequon);
            rankedSequons.add(rankedSequon);
            rank++;
        }

        return rankedSequons;
    }

}
```
