## Supplementary material for "Future Sequon Finder - A novel approach for predicting future N-linked glycosylation sequons on viral surface proteins": MegaInputProcessor.java

```
import java.util.ArrayList;
import java.util.List;


public class MegaInputProcessor {
    public static List<MegaSiteRate> processInput(String input) {
        List<MegaSiteRate> siteRates = new ArrayList<>();

        String[] lines = input.split("\n");
        for (String line : lines) {
            String[] parts = line.split(",");
            if (parts.length >= 2) {
                int position = Integer.parseInt(parts[0].trim());
                double rate = Double.parseDouble(parts[1].trim());
                rate = Math.round(rate * 10000.0) / 10000.0; // Round to 4 decimal places
                MegaSiteRate siteRate = new MegaSiteRate(position, rate);
                siteRates.add(siteRate);
            }
        }

        return siteRates;
    }
}
```
