## Supplementary material for "Future Sequon Finder - A novel approach for predicting future N-linked glycosylation sequons on viral surface proteins": MegaSiteRate.java

```
public class MegaSiteRate {
    private int positionIdentifier;
    private double mutationRate;

    public MegaSiteRate(int positionIdentifier, double mutationRate) {
        this.positionIdentifier = positionIdentifier;
        this.mutationRate = mutationRate;
    }

    public int getPositionIdentifier() {
        return positionIdentifier;
    }

    public double getMutationRate() {
        return mutationRate;
    }

}
```
