## Supplementary material for "Future Sequon Finder - A novel approach for predicting future N-linked glycosylation sequons on viral surface proteins": NearSequonScanner.java

```
import java.util.ArrayList;
import java.util.List;

class NearSequonScanner {
    public static List<String> findNearSequons(List<AminoAcid> aminoAcids, List<String> sequons) {
        List<String> nearSequonsList = new ArrayList<>();
        int nearSequonId = 1;

        for (int i = 0; i < aminoAcids.size() - 2; i++) {
            AminoAcid aa1 = getNextNonDeletionAA(aminoAcids, i);
            AminoAcid aa2 = getNextNonDeletionAA(aminoAcids, i + 1);
            AminoAcid aa3 = getNextNonDeletionAA(aminoAcids, i + 2);

            // Checks for perfect sequon match to eliminate from near-sequon consideration
            if (aa1.getAminoAcid().contains("N") && !aa2.getAminoAcid().contains("P") && (aa3.getAminoAcid().contains("S") || aa3.getAminoAcid().contains("T"))){
                continue;
            }
            // Check for N-P-S/T pattern
            if (aa1.getAminoAcid().contains("N") && aa2.getAminoAcid().contains("P") && (aa3.getAminoAcid().contains("S") || aa3.getAminoAcid().contains("T"))){
                String nearSequon = aa1.getAminoAcid() + aa2.getAminoAcid() + aa3.getAminoAcid();
                int aminoAcidPosition = i + 1;
                int nucleotidePosition = (((i+1) * 3)-3);
                String mismatchedCodon = mismatchedCodonAA2(nucleotidePosition, aminoAcids);
                String mismatchedAA = aa2.getAminoAcid();

                boolean singleNucleotideChange = true;

                boolean sameCharge = false;

                if (mismatchedAA.contains("G") || mismatchedAA.contains("A") || mismatchedAA.contains("V") || mismatchedAA.contains("C") || mismatchedAA.contains("L") || mismatchedAA.contains("I") || mismatchedAA.contains("M") || mismatchedAA.contains("W") || mismatchedAA.contains("F")){
                    sameCharge = true;
                }

                nearSequonsList.add("Near-Sequon " + nearSequonId + " at AA position " + aminoAcidPosition + ": " + nearSequon + ", Nucleotide position " + (nucleotidePosition + 1) + ": " + nucleotidePositionToNucleotides(nucleotidePosition, aminoAcids) + ", Mismatched codon (AA2 - " + mismatchedAA + "): " + mismatchedCodon + ", Single nucleotide change: " + singleNucleotideChange + ", Mismatched nucleotide in mismatched codon: NA" + ", Same charge & polarity: " + sameCharge);
                nearSequonId++;
            }
            // Check for _-!P-S/T pattern
            else if (!aa1.getAminoAcid().contains("N") && !aa2.getAminoAcid().contains("P") && (aa3.getAminoAcid().contains("S") || aa3.getAminoAcid().contains("T"))){
                String nearSequon = aa1.getAminoAcid() + aa2.getAminoAcid() + aa3.getAminoAcid();
                int aminoAcidPosition = i + 1;
                int nucleotidePosition = (((i+1) * 3)-3);
                int mismatchedCodonAAint = 1;
                String mismatchedCodon = mismatchedCodonAA1(nucleotidePosition, aminoAcids);
                String mismatchedAA = aa1.getAminoAcid();

                boolean singleNucleotideChange = false;
                int singleNucleotideChangePosition = isSingleNucleotideChange(mismatchedCodon, mismatchedCodonAAint);


                if (singleNucleotideChangePosition >= 0) {
                    singleNucleotideChange = true;
                } else {
                    singleNucleotideChangePosition = -2;
                }

                boolean sameCharge = false;

                if (mismatchedAA.contains("S") || mismatchedAA.contains("T") || mismatchedAA.contains(("Q"))){
                    sameCharge = true;
                }

                nearSequonsList.add("Near-Sequon " + nearSequonId + " at AA position " + aminoAcidPosition + ": " + nearSequon + ", Nucleotide position " + (nucleotidePosition + 1) + ": " + nucleotidePositionToNucleotides(nucleotidePosition, aminoAcids) + ", Mismatched codon (AA1 - " + mismatchedAA + "): " + mismatchedCodon + ", Single nucleotide change: " + singleNucleotideChange + ", Mismatched nucleotide in mismatched codon: " + (singleNucleotideChangePosition + 1) + ", Same charge & polarity: " + sameCharge);
                nearSequonId++;
            }
            // Check for N-!P-_ pattern
            else if (aa1.getAminoAcid().contains("N") && !aa2.getAminoAcid().contains("P") && !(aa3.getAminoAcid().contains("S") && !aa3.getAminoAcid().contains("T"))){
                String nearSequon = aa1.getAminoAcid() + aa2.getAminoAcid() + aa3.getAminoAcid();
                int aminoAcidPosition = i + 1;
                int nucleotidePosition = (((i+1) * 3)-3);
                int mismatchedCodonAAint = 3;
                String mismatchedCodon = mismatchedCodonAA3(nucleotidePosition, aminoAcids);
                String mismatchedAA = aa3.getAminoAcid();

                boolean singleNucleotideChange = false;
                int singleNucleotideChangePosition = isSingleNucleotideChange(mismatchedCodon, mismatchedCodonAAint);


                if (singleNucleotideChangePosition >= 0) {
                    singleNucleotideChange = true;
                } else {
                    singleNucleotideChangePosition = -2;
                }


                boolean sameCharge = false;

                if (mismatchedAA.contains("N") || mismatchedAA.contains(("Q"))){
                    sameCharge = true;
                }

                nearSequonsList.add("Near-Sequon " + nearSequonId + " at AA position " + aminoAcidPosition + ": " + nearSequon + ", Nucleotide position " + (nucleotidePosition + 1) + ": " + nucleotidePositionToNucleotides(nucleotidePosition, aminoAcids) + ", Mismatched codon (AA3 - " + mismatchedAA + "): " + mismatchedCodon + ", Single nucleotide change: " + singleNucleotideChange + ", Mismatched nucleotide in mismatched codon: " + (singleNucleotideChangePosition + 1) + ", Same charge & polarity: " + sameCharge);
                nearSequonId++;
            }
        }

        return nearSequonsList;
    }

    private static AminoAcid getNextNonDeletionAA(List<AminoAcid> aminoAcids, int start) {
        for (int i = start; i < aminoAcids.size(); i++) {
            if (!aminoAcids.get(i).getAminoAcid().equals("-")) {
                return aminoAcids.get(i);
            }
        }
        return null; // Allows for detection of near-sequons when deletions ("-" characters) are present in sequence
    }

    private static String nucleotidePositionToNucleotides(int position, List<AminoAcid> aminoAcids) {
        StringBuilder nucleotides = new StringBuilder();
        int nucleotideIndex = position;

        while (nucleotideIndex < position + 9 && nucleotideIndex < aminoAcids.size() * 3) {
            nucleotides.append(aminoAcids.get(nucleotideIndex / 3).getNucleotides().charAt(nucleotideIndex % 3));
            nucleotideIndex++;
        }
        return nucleotides.toString();
    }

    private static String mismatchedCodonAA1(int position, List<AminoAcid> aminoAcids) {
        StringBuilder nucleotides = new StringBuilder();
        int nucleotideIndex = position;

        while (nucleotideIndex < position + 3 && nucleotideIndex < aminoAcids.size() * 3) {
            nucleotides.append(aminoAcids.get(nucleotideIndex / 3).getNucleotides().charAt(nucleotideIndex % 3));
            nucleotideIndex++;
        }
        return nucleotides.toString();
    }

    private static String mismatchedCodonAA2(int position, List<AminoAcid> aminoAcids) {
        StringBuilder nucleotides = new StringBuilder();
        int nucleotideIndex = position + 3;

        while (nucleotideIndex < position + 6 && nucleotideIndex < (aminoAcids.size() * 3)) {
            nucleotides.append(aminoAcids.get(nucleotideIndex / 3).getNucleotides().charAt(nucleotideIndex % 3));
            nucleotideIndex++;
        }
        return nucleotides.toString();
    }

    private static String mismatchedCodonAA3(int position, List<AminoAcid> aminoAcids) {
        StringBuilder nucleotides = new StringBuilder();
        int nucleotideIndex = position + 6;

        while (nucleotideIndex < position + 9 && nucleotideIndex < ((aminoAcids.size()+1) * 3)) {
            nucleotides.append(aminoAcids.get(nucleotideIndex / 3).getNucleotides().charAt(nucleotideIndex % 3));
            nucleotideIndex++;
        }
        return nucleotides.toString();
    }

    private static int isSingleNucleotideChange(String mismatchedCodon, int mismatchedCodonAA) {
        String[] targetCodons;
        if (mismatchedCodonAA == 1) {
            targetCodons = new String[]{"AAT", "AAC"};
        } else if (mismatchedCodonAA == 3) {
            targetCodons = new String[]{"ACT", "ACC", "ACA", "TCT", "TCC", "TCA", "TCG", "ACG", "AGT", "AGC"};
        } else {
            return -2;
        }

        for (String targetCodon : targetCodons) {
            int index = indexOfSingleNucleotideDifference(mismatchedCodon, targetCodon);
            if (index != -1) {
                return index;
            }
        }

        return -1;
    }


    private static int indexOfSingleNucleotideDifference(String codon1, String codon2) {
        int differences = 0;
        int index = -1;
        for (int i = 0; i < codon1.length(); i++) {
            if (codon1.charAt(i) != codon2.charAt(i)) {
                differences++;
                index = i;
                if (differences > 1) {
                    return -1;
                }
            }
        }
        return differences == 1 ? index : -1;
    }

}
```
