## Supplementary material for "Future Sequon Finder - A novel approach for predicting future N-linked glycosylation sequons on viral surface proteins": NucleotideToAminoAcid.java

```
import java.util.ArrayList;
import java.util.List;


class NucleotideToAminoAcid {
    private static final String[] codons = {
            "TTT", "TTC", "TTA", "TTG", "CTT", "CTC", "CTA", "CTG",
            "ATT", "ATC", "ATA", "ATG", "GTT", "GTC", "GTA", "GTG",
            "TCT", "TCC", "TCA", "TCG", "CCT", "CCC", "CCA", "CCG",
            "ACT", "ACC", "ACA", "ACG", "GCT", "GCC", "GCA", "GCG",
            "TAT", "TAC", "TAA", "TAG", "CAT", "CAC", "CAA", "CAG",
            "AAT", "AAC", "AAA", "AAG", "GAT", "GAC", "GAA", "GAG",
            "TGT", "TGC", "TGA", "TGG", "CGT", "CGC", "CGA", "CGG",
            "AGT", "AGC", "AGA", "AGG", "GGT", "GGC", "GGA", "GGG",
            "---"
    };

    private static final String[] aminoAcids = {
            "F", "F", "L", "L", "L", "L", "L", "L",
            "I", "I", "I", "M", "V", "V", "V", "V",
            "S", "S", "S", "S", "P", "P", "P", "P",
            "T", "T", "T", "T", "A", "A", "A", "A",
            "Y", "Y", "_", "_", "H", "H", "Q", "Q",
            "N", "N", "K", "K", "D", "D", "E", "E",
            "C", "C", "_", "W", "R", "R", "R", "R",
            "S", "S", "R", "R", "G", "G", "G", "G",
            "-"
    };


    public static List<AminoAcid> convertToAminoAcids(String sequence) {
        List<AminoAcid> aminoAcidsList = new ArrayList<>();
        int aaOrder = 1;
        int nucleotideOrder = 1;

        for (int i = 0; i < sequence.length(); i += 3) {
            if (i + 2 >= sequence.length()) {
                break;
            }

            String codon = sequence.substring(i, Math.min(i + 3, sequence.length()));

            if (codon.contains("-")) {
                aminoAcidsList.add(new AminoAcid("-", codon, String.valueOf(nucleotideOrder), aaOrder));
                aaOrder++;
                String combinedCodon = "";
                int j = i;
                while (combinedCodon.length() < 3 && j < sequence.length()) {
                    combinedCodon += sequence.charAt(j) != '-' ? sequence.charAt(j) : "";
                    j++;
                }

                if (combinedCodon.length() >= 3) {
                    combinedCodon = combinedCodon.substring(0, 3);
                    int aminoAcidIndex = findAminoAcidIndex(combinedCodon);
                    if (aminoAcidIndex != -1) {
                        String aminoAcidCode = aminoAcids[aminoAcidIndex];
                        aminoAcidsList.add(new AminoAcid(aminoAcidCode, combinedCodon, nucleotideOrder + "-" + (j - 1), aaOrder));
                    }
                    i = j - 3;
                }
                nucleotideOrder += j - i;
                aaOrder++;
            } else {
                int aminoAcidIndex = findAminoAcidIndex(codon);
                if (aminoAcidIndex != -1) {
                    String aminoAcidCode = aminoAcids[aminoAcidIndex];
                    aminoAcidsList.add(new AminoAcid(aminoAcidCode, codon, String.valueOf(nucleotideOrder), aaOrder));
                }
                nucleotideOrder += 3;
                aaOrder++;
            }
        }

        return aminoAcidsList;
    }

    private static int findAminoAcidIndex(String codon) {
        for (int j = 0; j < codons.length; j++) {
            if (codons[j].equals(codon)) {
                return j;
            }
        }
        return -1;
    }
}
```
