## Supplementary material for "Future Sequon Finder - A novel approach for predicting future N-linked glycosylation sequons on viral surface proteins": SequonScanner.java

```
import java.util.ArrayList;
import java.util.List;

class SequonScanner {
    public static List<String> findSequons(List<AminoAcid> aminoAcids) {
        List<String> sequonsList = new ArrayList<>();
        int sequonId = 1;

        for (int i = 0; i < aminoAcids.size() - 2; i++) {
            AminoAcid aa1 = aminoAcids.get(i);
            AminoAcid aa2 = getNextNonDeletionAA(aminoAcids, i + 1);
            AminoAcid aa3 = getNextNonDeletionAA(aminoAcids, i + 2);

            if (aa1.getAminoAcid().equals("N") && !aa2.getAminoAcid().equals("P") && (aa3.getAminoAcid().equals("S") || aa3.getAminoAcid().equals("T"))) {
                sequonsList.add("Sequon " + sequonId + " at AA position " + aa1.getAminoAcidOrder() + ": " + aa1.getAminoAcid() + aa2.getAminoAcid() + aa3.getAminoAcid());
                sequonId++;
            }
        }

        return sequonsList;
    }

    private static AminoAcid getNextNonDeletionAA(List<AminoAcid> aminoAcids, int start) {
        for (int i = start; i < aminoAcids.size(); i++) {
            if (!aminoAcids.get(i).getAminoAcid().equals("-")) {
                return aminoAcids.get(i);
            }
        }
        return null; // Allows for detection of sequons when deletions ("-" characters) are present in sequence
    }
}
```
