## Supplementary material for "Future Sequon Finder - A novel approach for predicting future N-linked glycosylation sequons on viral surface proteins": Main.java

```
import javax.swing.*;
import java.awt.*;
import java.util.concurrent.atomic.AtomicInteger;
import java.util.regex.*;
import javax.swing.border.Border;
import java.util.HashMap;
import java.util.Map;
import java.util.HashSet;
import java.util.Set;
import java.util.ArrayList;
import java.util.Collections;
import java.util.List;
import java.util.Random;

public class Main {

    // To use the program simply input a nucleotide FASTA file and the program will return an AA FASTA file. It has some selection methods to get rid of unwanted sequences and has been tuned for HA proteins. If you are using another kind of protein, modify the code according to your needs.
    public static void main(String[] args) {
        JFrame frame = new JFrame("Influenza Fasta Sorter");
        frame.setDefaultCloseOperation(JFrame.EXIT_ON_CLOSE);
        frame.setLayout(new BorderLayout(5, 10));
        frame.setSize(1400, 1000);

        Border padding = BorderFactory.createEmptyBorder(10, 10, 10, 10);
        ((JComponent) frame.getContentPane()).setBorder(padding);

        JTextArea outputTextArea = new JTextArea("Please provide inputs and press 'Run'", 20, 20);
        outputTextArea.setEditable(false);
        JScrollPane outputScrollPane = new JScrollPane(outputTextArea);
        frame.add(outputScrollPane, BorderLayout.NORTH);

        JPanel fastaPanel = new JPanel(new BorderLayout());
        fastaPanel.add(new JLabel("Input FASTA sequences below:"), BorderLayout.NORTH);
        JTextArea fastaInputTextArea = new JTextArea(10, 20);
        fastaInputTextArea.setLineWrap(true);
        JScrollPane fastaInputScrollPane = new JScrollPane(fastaInputTextArea);
        fastaPanel.add(fastaInputScrollPane, BorderLayout.CENTER);

        JPanel cutoffYearPanel = new JPanel(new FlowLayout(FlowLayout.LEFT));
        cutoffYearPanel.add(new JLabel("Input cutoff year:"));
        JTextField cutoffYearField = new JTextField(20);
        cutoffYearPanel.add(cutoffYearField);

        JPanel maxSequencesPanel = new JPanel(new FlowLayout(FlowLayout.LEFT));
        maxSequencesPanel.add(new JLabel("Maximum sequences per year:"));
        JTextField maxSequencesField = new JTextField(20);
        maxSequencesPanel.add(maxSequencesField);

        // Show summary information checkbox
        JCheckBox showSummaryCheckbox = new JCheckBox("Show summary information:", true);
        JPanel showSummaryPanel = new JPanel(new FlowLayout(FlowLayout.LEFT));
        showSummaryPanel.add(showSummaryCheckbox);

        JPanel centerPanel = new JPanel();
        centerPanel.setLayout(new BoxLayout(centerPanel, BoxLayout.Y_AXIS));
        centerPanel.add(fastaPanel);
        centerPanel.add(cutoffYearPanel);
        centerPanel.add(maxSequencesPanel);
        centerPanel.add(showSummaryPanel);
        frame.add(centerPanel, BorderLayout.CENTER);

        JPanel bottomPanel = new JPanel(new FlowLayout(FlowLayout.RIGHT));
        JButton resetButton = new JButton("Reset");
        resetButton.addActionListener(e -> {
            fastaInputTextArea.setText("");
            cutoffYearField.setText("");
            maxSequencesField.setText("");
            outputTextArea.setText("Please provide inputs and press 'Run'");
        });
        bottomPanel.add(resetButton);

        JButton runButton = new JButton("Run");
        runButton.addActionListener(e -> {
            boolean showSummary = showSummaryCheckbox.isSelected();
            processInputs(fastaInputTextArea.getText(), cutoffYearField.getText(), maxSequencesField.getText(), outputTextArea, showSummary); // Pass showSummary to processInputs
        });
        bottomPanel.add(runButton);

        frame.add(bottomPanel, BorderLayout.SOUTH);

        frame.setVisible(true);
    }

    private static void processInputs(String fastaData, String cutoffYearStr, String maxSequencesField, JTextArea outputTextArea, boolean showSummary) {
        outputTextArea.setText("");

        int cutoffYear;
        try {
            cutoffYear = Integer.parseInt(cutoffYearStr);
        } catch (NumberFormatException e) {
            outputTextArea.append("Invalid cutoff year.\n");
            return;
        }

        int maxPerYear = maxSequencesField.trim().isEmpty() ? Integer.MAX_VALUE : Integer.parseInt(maxSequencesField);

        Map<Integer, List<String>> sequencesByYear = new HashMap<>();
        int totalSequencesEntered = 0;
        AtomicInteger totalSequencesDisplayed = new AtomicInteger();
        int totalExcludedByYear = 0;
        int totalExcludedByCharacters = 0;
        int duplicatesRemoved = 0;
        AtomicInteger totalExcludedByMaxPerYear = new AtomicInteger();
        Set<String> uniqueNames = new HashSet<>();

        String[] entries = fastaData.split(">");
        for (String entry : entries) {
            if (entry.trim().isEmpty()) continue;
            totalSequencesEntered++;

            String[] lines = entry.split("\n", 2);
            if (lines.length < 2) continue;

            String header = lines[0];
            String sequence = lines[1].replaceAll("\\s+", "").toUpperCase();

            String name = extractName(header).replace("-", "");
            int year = extractYearFromName(name);

            if (!uniqueNames.add(name)) {
                duplicatesRemoved++;
                continue;
            }

            if (year == -1 || year > cutoffYear) {
                totalExcludedByYear++;
                continue;
            }

            if (!sequence.matches("[ATGC]+")) {
                totalExcludedByCharacters++;
                continue;
            }

            if (sequence.length() < 1695) {
                totalExcludedByCharacters++;
                continue;
            }

            Matcher m = Pattern.compile("ATG").matcher(sequence);
            if (m.find()) {
                sequence = sequence.substring(m.start());
            } else {
                totalExcludedByCharacters++;
                continue;
            }

            // Ensure all sequences are nearly identical in length (No H1N1 should exceed 1701, no H3N2 should exceed 1704)
            if (sequence.length() > 1704) {
                sequence = sequence.substring(0, 1704);
            }

            String aminoAcidSequence = NucleotideToAminoAcidConverter.translateToAminoAcids(sequence);

            // Trim to the first stop codon
            int stopCodonIndex = aminoAcidSequence.indexOf("_");
            if (stopCodonIndex != -1) {
                aminoAcidSequence = aminoAcidSequence.substring(0, stopCodonIndex);
            }

            if (aminoAcidSequence.length() < 565 || aminoAcidSequence.length() >= 567) {
                totalExcludedByCharacters++;
                continue;
            }


            String fastaEntry = ">" + name + "\n" + aminoAcidSequence;
            sequencesByYear.computeIfAbsent(year, k -> new ArrayList<>()).add(fastaEntry);
        }

        Random random = new Random();
        sequencesByYear.forEach((year, sequences) -> {
            if (sequences.size() > maxPerYear) {
                Collections.shuffle(sequences, random);
                List<String> selectedSequences = new ArrayList<>(sequences.subList(0, maxPerYear));
                totalExcludedByMaxPerYear.addAndGet(sequences.size() - maxPerYear);
                sequences.clear();
                sequences.addAll(selectedSequences); // Update the list with selected sequences
            }
            sequences.forEach(sequence -> {
                outputTextArea.append(sequence + "\n");
                totalSequencesDisplayed.getAndIncrement();
            });
        });


        if (showSummary) {
            // Append summary information
            outputTextArea.append("\nTotal sequences entered: " + totalSequencesEntered);
            outputTextArea.append("\nTotal sequences displayed: " + totalSequencesDisplayed);
            outputTextArea.append("\nTotal sequences excluded due to cutoff year: " + totalExcludedByYear);
            outputTextArea.append("\nTotal sequences excluded due to invalid formatting: " + totalExcludedByCharacters);
            outputTextArea.append("\nDuplicates removed: " + duplicatesRemoved + "\n");
            outputTextArea.append("\nSequences per year:\n");
            sequencesByYear.forEach((year, sequences) -> outputTextArea.append(year + ": " + sequences.size() + " sequences\n"));
        }
    }

    private static String extractName(String header) {
        Pattern namePattern = Pattern.compile("\\((.*?)\\((H1N1|H3N2)\\)\\)");
        Matcher matcher = namePattern.matcher(header);
        if (matcher.find()) {
            return (matcher.group(1) + "(" + matcher.group(2) + ")").replace("-", "");
        }
        return "Name not found";
    }

    private static int extractYearFromName(String name) {
        try {
            String[] parts = name.split("/");
            String lastPart = parts[parts.length - 1];
            // Use a regular expression to find the subtype (either H1N1 or H3N2) within the last part of the name. I needed this to determine what year my strains were collected.
            Pattern pattern = Pattern.compile("(H1N1|H3N2)");
            Matcher matcher = pattern.matcher(lastPart);
            String yearStr = "";
            if (matcher.find()) {
                // Extract substring up to the found subtype
                yearStr = lastPart.substring(0, lastPart.indexOf(matcher.group()));
            }
            String yearDigits = yearStr.replaceAll("\\D+", "");
            int year;
            if (yearDigits.length() == 2) {
                int yearInt = Integer.parseInt(yearDigits);
                // Check if the year is less than 24, then prefix with "20", else "19"
                if (yearInt < 24) {
                    year = Integer.parseInt("20" + yearDigits);
                } else {
                    year = Integer.parseInt("19" + yearDigits);
                }
            } else if (yearDigits.length() == 4) {
                year = Integer.parseInt(yearDigits);
            } else {
                throw new Exception("Invalid year format");
            }
            return year;
        } catch (Exception e) {
            System.out.println("Error extracting year from name: " + name + ". Error: " + e.getMessage());
            return -1;
        }
    }

}
```
