## Supplementary material for "Future Sequon Finder - A novel approach for predicting future N-linked glycosylation sequons on viral surface proteins": NucleotideToAminoAcidConverter.java

```
public class NucleotideToAminoAcidConverter {
    private static final String[] codons = {
            "TTT", "TTC", "TTA", "TTG", "CTT", "CTC", "CTA", "CTG",
            "ATT", "ATC", "ATA", "ATG", "GTT", "GTC", "GTA", "GTG",
            "TCT", "TCC", "TCA", "TCG", "CCT", "CCC", "CCA", "CCG",
            "ACT", "ACC", "ACA", "ACG", "GCT", "GCC", "GCA", "GCG",
            "TAT", "TAC", "TAA", "TAG", "CAT", "CAC", "CAA", "CAG",
            "AAT", "AAC", "AAA", "AAG", "GAT", "GAC", "GAA", "GAG",
            "TGT", "TGC", "TGA", "TGG", "CGT", "CGC", "CGA", "CGG",
            "AGT", "AGC", "AGA", "AGG", "GGT", "GGC", "GGA", "GGG",
    };

    private static final String[] aminoAcids = {
            "F", "F", "L", "L", "L", "L", "L", "L",
            "I", "I", "I", "M", "V", "V", "V", "V",
            "S", "S", "S", "S", "P", "P", "P", "P",
            "T", "T", "T", "T", "A", "A", "A", "A",
            "Y", "Y", "_", "_", "H", "H", "Q", "Q",
            "N", "N", "K", "K", "D", "D", "E", "E",
            "C", "C", "_", "W", "R", "R", "R", "R",
            "S", "S", "R", "R", "G", "G", "G", "G",
    };

    public static String translateToAminoAcids(String nucleotideSequence) {
        StringBuilder aminoAcidSequence = new StringBuilder();
        for (int i = 0; i + 2 < nucleotideSequence.length(); i += 3) {
            String codon = nucleotideSequence.substring(i, i + 3).toUpperCase();
            int index = -1;
            for (int j = 0; j < codons.length; j++) {
                if (codons[j].equals(codon)) {
                    index = j;
                    break;
                }
            }
            if (index != -1) {
                aminoAcidSequence.append(aminoAcids[index]);
            } else {
                aminoAcidSequence.append("X");
            }
        }
        return aminoAcidSequence.toString();
    }
}
```
