## Supplementary material for "Future Sequon Finder - A novel approach for predicting future N-linked glycosylation sequons on viral surface proteins": Main.java

```
import javax.swing.*;
import java.awt.*;
import java.awt.event.ActionEvent;
import java.awt.event.ActionListener;
import javax.swing.border.Border;

class Main {
    public static void main(String[] args) {
        JFrame frame = new JFrame("Multiple Sequon Finder");
        frame.setDefaultCloseOperation(JFrame.EXIT_ON_CLOSE);
        frame.setSize(800, 600);

        frame.setLayout(new BorderLayout());

        JTextArea outputArea = new JTextArea("Welcome to Multiple Sequon Finder! Please input your aligned amino acid sequences in .meg format below.");
        outputArea.setEditable(false);
        outputArea.setRows(10);
        outputArea.setFont(new Font("Monospaced", Font.PLAIN, 12));  // Set the font to Monospaced
        JScrollPane outputScrollPane = new JScrollPane(outputArea);
        frame.add(outputScrollPane, BorderLayout.CENTER);

        JPanel southPanel = new JPanel(new BorderLayout());

        JTextArea inputTextArea = new JTextArea(5, 20);
        inputTextArea.setLineWrap(true);
        inputTextArea.setWrapStyleWord(true);
        inputTextArea.setFont(new Font("Monospaced", Font.PLAIN, 12));  // Optionally, set the font for input area as well
        JScrollPane inputScrollPane = new JScrollPane(inputTextArea);
        JPanel inputPanel = new JPanel(new BorderLayout());
        inputPanel.add(new JLabel("Input aligned sequences here (.MEG format):"), BorderLayout.NORTH);
        inputPanel.add(inputScrollPane, BorderLayout.CENTER);
        southPanel.add(inputPanel, BorderLayout.CENTER);

        JPanel buttonPanel = new JPanel(new FlowLayout(FlowLayout.RIGHT));
        JButton submitButton = new JButton("Submit");
        JButton resetButton = new JButton("Reset");

        submitButton.addActionListener(new ActionListener() {
            @Override
            public void actionPerformed(ActionEvent e) {
                String[] sequences = inputTextArea.getText().split("#");
                SequonScanner scanner = new SequonScanner();
                for (String seq : sequences) {
                    if (!seq.trim().isEmpty() && !seq.contains("mega") && !seq.contains("!Title") && !seq.contains("!Format")) {
                        String[] lines = seq.trim().split("\\s+", 2);
                        if (lines.length > 1) {
                            scanner.scan(lines[0].trim(), lines[1].trim());
                        }
                    }
                }
                outputArea.setText(scanner.getFormattedResults());
            }
        });

        resetButton.addActionListener(new ActionListener() {
            @Override
            public void actionPerformed(ActionEvent e) {
                inputTextArea.setText("");
                outputArea.setText("Welcome to Multiple Sequon Finder! Please input your aligned amino acid sequences in .meg format below.");
            }
        });

        buttonPanel.add(submitButton);
        buttonPanel.add(resetButton);
        southPanel.add(buttonPanel, BorderLayout.SOUTH);
        frame.add(southPanel, BorderLayout.SOUTH);
        frame.setVisible(true);
    }
}
```
