## Supplementary material for "Future Sequon Finder - A novel approach for predicting future N-linked glycosylation sequons on viral surface proteins": SequonScanner.java

```
import java.util.*;
import java.util.stream.Collectors;

public class SequonScanner {
    private Map<Integer, Set<String>> sequonSites = new LinkedHashMap<>();
    private int totalStrains = 0;  // To keep track of the total number of strains entered

    public void scan(String name, String sequence) {
        totalStrains++;  // Increment the total strains count for each scanned strain
        sequence = sequence.replaceAll("\\s+", ""); // Remove all whitespaces
        int actualIndex = 0; // Index without considering "-"

        // Loop through the sequence, adjusting for sequons
        for (int i = 0; i < sequence.length(); i++) {
            if (sequence.charAt(i) == '-') {
                actualIndex++;
                continue;
            }
            if (i + 2 < sequence.length() &&
                    sequence.charAt(i) == 'N' &&
                    sequence.charAt(i + 1) != 'P' &&
                    (sequence.charAt(i + 2) == 'S' || sequence.charAt(i + 2) == 'T') &&
                    sequence.charAt(i + 1) != '-' && // Ensure the sequon isn't interrupted by dashes
                    sequence.charAt(i + 2) != '-') {
                sequonSites.computeIfAbsent(actualIndex + 1, k -> new HashSet<>()).add(name);
            }
            actualIndex++;
        }
    }

    public String getFormattedResults() {
        StringBuilder sb = new StringBuilder();
        sb.append("Total number of strains entered: ").append(totalStrains).append("\n\n");

        List<Integer> sortedKeys = new ArrayList<>(sequonSites.keySet());
        Collections.sort(sortedKeys);

        // Determine maximum widths for table columns
        int maxSequonLabelWidth = sortedKeys.stream()
                .mapToInt(key -> Integer.toString(key).length())
                .max()
                .orElse(0) + 15; // Increased padding for better spacing

        int maxTotalStrainsWidth = sequonSites.values().stream()
                .mapToInt(set -> Integer.toString(set.size()).length())
                .max()
                .orElse(0) + 15; // Increased padding for better spacing

        int maxPercentageWidth = 15; // Increased fixed width for percentage to allow for more spacing

        // Construct table headers
        sb.append(String.format("%-" + maxSequonLabelWidth + "s", "Sequon site"))
                .append(String.format("%-" + maxTotalStrainsWidth + "s", "Total strains"))
                .append(String.format("%-" + maxPercentageWidth + "s", "Percentage"))
                .append("First strain\n");

        // Create a separator line
        int totalWidth = maxSequonLabelWidth + maxTotalStrainsWidth + maxPercentageWidth + "First strain".length();
        sb.append("-".repeat(totalWidth)).append("\n");

        // Fill table rows
        for (int key : sortedKeys) {
            Set<String> names = sequonSites.get(key);
            List<String> sortedNames = new ArrayList<>(names);
            Collections.sort(sortedNames, (o1, o2) -> extractYear(o1) - extractYear(o2));
            String oldestSequence = sortedNames.stream()
                    .filter(n -> extractYear(n) > 0)
                    .findFirst()
                    .orElse("No valid sequences found");

            int sequonCount = names.size();
            double percentage = 100.0 * sequonCount / totalStrains;

            sb.append(String.format("%-" + maxSequonLabelWidth + "s", key))  // Removed "Sequon site"
                    .append(String.format("%-" + maxTotalStrainsWidth + "s", sequonCount))
                    .append(String.format("%-" + maxPercentageWidth + "s", String.format("%.2f%%", percentage)))
                    .append(oldestSequence).append("\n");
        }
        return sb.toString();
    }


    private int extractYear(String name) {
        try {
            int startIndex = name.lastIndexOf("/") + 1;
            int endIndex = name.indexOf("(", startIndex);
            if (startIndex < 1 || endIndex == -1 || startIndex >= endIndex) {
                return 0;  // Invalid indices for year extraction
            }
            String yearPart = name.substring(startIndex, endIndex).trim();
            if (!yearPart.matches("\\d{4}")) {
                return 0;  // Not a valid year format
            }
            return Integer.parseInt(yearPart);
        } catch (NumberFormatException e) {
            return 0;  // Handle cases where the year is not properly formatted
        }
    }
}
```
